# Antibody recycling via FcRn drives atherosclerotic plaque vulnerability

**DOI:** 10.64898/2026.03.08.710352

**Authors:** Shiying Lin, Justine Deroissart, Yinda Yu, Yanzhao Wu, Martina B Lorey, Lena Steiger, Xintong Jiang, Glykeria Karadimou, Stephen G Malin, Katariina Öörni, Ulf Hedin, Christoph J Binder, Anton Gisterå

## Abstract

Atherosclerotic plaques accumulate low-density lipoprotein (LDL) together with antibodies targeting LDL and its apolipoprotein B (apoB) component. Given the association between IgG and plaque vulnerability, we hypothesized that apoB-specific immune complexes actively promote plaque destabilization. Using immunohistochemistry in carotid endarterectomy specimens, we quantified antibody deposition across morphologically defined plaque regions, and measured apoB reactivity and immune complex levels in matched plaque and plasma samples.

IgG deposition was strongly associated with thin fibrous caps, reduced collagen content, and higher overall plaque vulnerability. Symptomatic patients exhibited increased apoB-specific IgG and reduced apoB-IgG immune complexes within plaques, indicating enhanced IgG recycling and heightened inflammatory activity. The neonatal Fc receptor (FcRn) was predominantly expressed by CD163^+^ macrophages, and mediated antibody recycling, LDL uptake, and production of tumor necrosis factor (TNF) and matrix metalloproteinase-9 (MMP-9) in vitro. Plaque FcRn expression increased with age and correlated with mediators of vulnerability, including collagen-degrading enzymes and pro-inflammatory cytokines. Ex vivo treatment of human plaques with a clinically used FcRn-blocking monoclonal antibody reduced IgG recycling and suppressed TNF and MMP-9 production.

These findings identify FcRn-dependent antibody recycling as a contributor to inflammatory plaque vulnerability and highlight FcRn as a potential therapeutic target in atherosclerosis.

## Introduction

Atherosclerosis is a chronic inflammatory disease driven by the arterial retention of apolipoprotein (apo) B-containing lipoproteins, which initiates and sustains plaque formation [1, 2]. Although autoimmune responses to apoB have been described, their precise contribution to disease progression remains unresolved [3]. As plaques mature, structural destabilization through erosion or rupture can trigger thrombosis, leading to myocardial infarction or ischemic stroke.

Genetic evidence establishes a direct, causal relationship between plasma apoB levels and atherosclerotic cardiovascular risk [4]. ApoB-containing lipoproteins, particularly those in the low-density lipoprotein (LDL) fraction, cross the endothelium and accumulate subendothelially, where they undergo enzymatic and oxidative modifications. These modified LDL particles, including apoB-derived peptides and cholesterol crystals, activate both innate and adaptive immune pathways [5–7]. In peripheral lymphoid organs, LDL-reactive B cells receive help from apoB-specific T cells, undergo class switching, and generate high-affinity IgG antibodies. Both antibodies and apoB-reactive T cells subsequently accumulate within atherosclerotic lesions rich in retained LDL [3].

Consequently, atherosclerotic plaques contain abundant immunoglobulins, including IgG, IgA, and IgM [8]. Although early metabolic-labeling studies suggested that some antibodies may be synthesized within the plaque, the overall contribution of local production remains unclear given the scarcity of intraplaque B cells [9, 10]. Instead, antibody responses are thought to originate primarily in secondary or tertiary lymphoid organs, with B cells trafficking through the adventitia and draining lymph nodes to sustain ongoing antigen-specific immunity [11, 12].

Oxidative modification of LDL generates malondialdehyde-protein adducts and oxidized phospholipids, which act as immunogenic neoepitopes [13]. Both mice and humans mount antibody responses against these oxidation-specific epitopes; notably, IgM can neutralize them and inhibit macrophage uptake and foam-cell formation [14, 15]. Because both IgM and IgG also recognize native apoB, apoB-specific reactivity provides a broad yet selective measure of LDL-directed humoral immunity [16].

IgG-LDL immune complexes have been reported to exert context-dependent effects. In some settings, they facilitate lipoprotein clearance and dampen inflammation via inhibitory Fc receptor engagement [17–20]. In others, they instead amplify macrophage activation and inflammatory signaling [21]. This variability underscores the need for detailed plaque-level characterization to clarify how immune complex biology shapes local immune responses within atherosclerotic lesions.

Here, we sought to determine how apoB-containing immune complexes influence plaque vulnerability. Prior histopathological reports have described IgM enrichment near lipid cores of disrupted, high-risk plaques in fatal myocardial infarction cases [22]. Focusing on IgG, given its stronger links to pro-inflammatory activity, we found that human carotid plaques contain substantial IgG, with apoB-specific IgG significantly enriched in symptomatic patients. FcRn-expressing macrophages displayed increased uptake of apoB immune complexes, consistent with enhanced FcRn-dependent recycling, and FcRn blockade reversed this phenotype. These findings identify FcRn-mediated handling of apoB immune complexes as a driver and potential therapeutic target of plaque destabilization.

## Materials and Methods

### Carotid Endarterectomy Cohort and Sample Acquisition

The Biobank of Karolinska Endarterectomies (BiKE) includes patients undergoing carotid endarterectomy for symptomatic or asymptomatic high-grade carotid stenosis (>50% as defined by The North American Symptomatic Carotid Endarterectomy Trial; Table S1) [23, 24]. Carotid plaques were collected using previously described protocols, and peripheral ethylenediaminetetraacetic acid plasma was obtained before the procedure [25]. Non-diseased iliac arteries from organ donors served as controls [26]. Carotid plaque gene expression was profiled using Affymetrix microarrays (GSE21545).

All procedures were approved by the Stockholm Ethics Committee, adhered to the Declaration of Helsinki, and included written informed consent. Single-cell RNA-sequencing datasets (GSE131778, GSE155512, GSE253904, GSE159677, and DOI 10.5281/zenodo.6032098) were integrated for comparative analyses, and single-cell assay for transposase-accessible chromatin (ATAC) sequencing data were accessed through the PlaqView portal [27–32]. Macrophage subsets in murine atherosclerotic arteries were examined using single-cell RNA-sequencing data from *Ldlr*^-/-^ mice (GSE155513) [28]. Microdissected macrophage-rich regions from ruptured and stable human plaques were analyzed using Affymetrix microarrays (GSE41571) [33].

### Immunohistochemistry and Plaque Morphological Assessment

Five-µm paraffin sections of human carotid plaques were fixed in 4% phosphate-buffered formaldehyde, deparaffinized, and rehydrated through graded ethanol. Antigen retrieval was performed by high-pressure boiling in Diva Decloaker buffer (pH 6.0, Biocare Medical). Endogenous biotin was blocked using an avidin-biotin kit (Vector Labs), followed by incubation in 5% normal horse serum in tris-buffered saline with 0.1% Tween 20 (TBST). Primary antibodies (Table S2) were applied overnight at 4°C, and biotinylated secondary antibodies were added for 30 minutes at room temperature. Detection was performed using Avidin-Biotin-Complex alkaline phosphatase (Vector Labs) and Warp Red (Biocare Medical), with hematoxylin QS used for counterstaining.

For immunofluorescence, acetone-fixed cryosections were stained with fluorescent secondary antibodies for 60 minutes at room temperature, followed by 4′,6-diamidino-2-phenylindole nuclear staining (D1306, Invitrogen) and mounting in Fluorescent Mounting Medium (Dako). Images were acquired using a Nikon Eclipse Ti2 confocal microscope.

For collagen quantification, deparaffinized sections were stained with 0.1% Picrosirius Red (Histolab) and rinsed in 0.5% acetic acid before air-drying. These sections were used to determine minimal cap thickness and delineate cap, core, and shoulder regions. Calcification was assessed using 2% Alizarin Red (Merck), iron deposition with Perls staining (04-180807, Bio-Optica), and acidic tissue components with 1% Toluidine Blue.

Following dehydration in graded ethanol and xylene, sections were mounted with Pertex. Whole-slide imaging was performed using a Hamamatsu Nanozoomer or Olympus Slideview VS200. Two blinded operators quantified staining as the percentage of positive area relative to total plaque area using Leica Qwin or QuPath, with high inter-observer agreement (r > 0.9). A vulnerability index was calculated by summing ranks of stability features (cap thickness, collagen content, smooth muscle cell staining) and dividing by the summed ranks of vulnerability features (hemorrhage, necrotic core size, macrophage staining).

Pre-operative computed tomography angiography was evaluated using semiautomated plaque-morphology workflows to derive minimal cap thickness, lipid-rich necrotic core, and intraplaque hemorrhage from tissue-composition maps on cross-sectional carotid reconstructions [34]. A computed tomography-based vulnerability index was then calculated by ranking cap thickness and dividing this rank by the combined ranks of lipid-rich necrotic core and intraplaque hemorrhage.

### Plaque Protein Extraction and Immunoassays

Frozen carotid plaques were rinsed in cold phosphate-buffered saline, minced, and homogenized in radioimmunoprecipitation buffer containing protease inhibitors (Promega) using 1.4 mm ceramic beads (Precellys) in a Tissue Lyzer (Qiagen) for four 2-minute cycles at 30 Hz. Homogenates were briefly centrifuged at 2,000 g to remove foam and concentrated to 500 µl using 50 kDa molecular-weight cutoff filters (Thermo Fisher). Total protein content was quantified using the Pierce bicinchoninic acid protein assay kit (Thermo Fisher).

Antibody isotype levels were measured by enzyme-linked immunosorbent assay (ELISA; antibodies listed in Table S3). Plates were coated overnight at 4°C with capture antibodies, blocked with 1% bovine serum albumin in TBS, and incubated with plaque extracts or plasma (diluted 1:25-1:80,000) overnight at 4°C. Detection antibodies were applied for 2 hours at room temperature, and chemiluminescence was developed using Lumiphos Plus (Lumigen) and read on a BMG Labtech microplate reader.

ApoB reactivity was assessed by coating plates with 10 μg/ml of native apoB100 (318630, MyBioSource) overnight at 4°C and blocking with 1% gelatin. Plasma and plaque samples were added at final concentrations of 1 µg/ml IgG or 0.1 µg/ml IgM in 0.1% gelatin-TBS. Isotypes (IgM, IgG1-4) were detected using biotinylated anti-human antibodies, streptavidin-alkaline phosphatase, and chemiluminescent substrate.

For detection of apoB-IgG immune complexes, plates were coated with polyclonal anti-apoB and incubated with plasma (1:2,600) or plaque extracts (1:300), followed by alkaline phosphatase-conjugated detection antibodies. ApoB and matrix metalloproteinase-9 (MMP-9) concentrations were measured using commercial ELISA kits (3715-1H-6, Mabtech; KE00298, Proteintech).

### Immunoglobulin Isolation and Macrophage Experiments In vitro

Human IgG was isolated from plaque protein extracts, and mouse anti-LDL antibodies were purified from plasma of apoB-reactive T-cell transfer experiments using the BT3 TCR-transgenic strain and human *APOB100*-transgenic *Ldlr*^tm1Her^ (HuBL, European mutant mouse archive 09689) mice. All animal procedures were approved by the Stockholm Regional Animal Ethics Board. Total human and mouse IgG was purified by protein G affinity (Thermo Scientific), dialyzed against phosphate-buffered saline overnight at 4°C, and sterile-filtered through 0.22 µm polyvinylidene fluoride filters (Millipore).

For macrophage stimulation assays, plates were coated with 10 µg/ml oxidized (ox)LDL (Thermo Fisher) overnight at 37°C and washed before forming immune complexes by incubating with 20 µg/ml isolated IgG overnight at 37°C. RAW 264.7 macrophages (ATCC TIB-71) were added at 80% confluency in Roswell Park Memorial Institute (RPMI) 1640 containing 1% fetal bovine serum and 2 mM L-glutamine. *Fcgrt* knockdown was achieved using Lipofectamine 3000 and target-specific siRNAs (Table S4). After 24 hours, supernatants were collected for total IgG and tumor necrosis factor (TNF) ELISA (DY410, R&D Systems).

For human macrophage experiments, THP-1 cells (ATCC TIB-202) were differentiated with 100 ng/ml phorbol 12-myristate 13-acetate for 48 hours. Immune complexes were generated using 20 µg/ml isolated human plaque IgG. Cells were treated with 10 µg/ml rozanolixizumab (anti-human FcRn; HY-P9979, MedChemExpress) or an IgG4 isotype control (HY-P99003, MedChemExpress). After 24 hours, supernatants were analyzed for IgG1-3 and TNF by ELISA (DY210, R&D Systems).

Cytotoxicity was assessed using the CyQUANT LDH assay (C20300, Invitrogen). Cells were harvested for protein and RNA extraction. RNA was isolated using the mirVana kit (Invitrogen), and 250 ng RNA per sample was reverse-transcribed using the TaqMan MicroRNA Reverse Transcription Kit. qPCR was performed using assay-on-demand probes (Table S4), with expression normalized to Hprt or PPIA using the ΔΔCt method.

For immunoblotting, 20 µg protein lysates were resolved on 10% Mini-PROTEAN TGX gels (Bio-Rad), transferred to methanol-activated polyvinylidene difluoride membrane, blocked in 5% milk in TBST, and incubated with primary antibodies overnight at 4°C. HRP-conjugated secondary antibodies were applied for 1 hour, and signals were detected with SuperSignal West Femto (ThermoFisher) on a Bio-Rad ChemiDoc MP system. Membranes were stripped using Millipore stripping buffer.

Lipoprotein uptake assays used ATTO594-labeled LDL. Plates were coated with labeled LDL, and LDL-IgG immune complexes were formed before adding macrophages. After 24-40 hours at 37°C, single-cell suspensions were analyzed by flow cytometry (Cytek NL-3000). All in vitro experiments were performed at least twice.

### Ex vivo Stimulation of Human Carotid Plaque Tissue

Fresh carotid plaques from symptomatic patients were collected immediately after endarterectomy and transferred into sterile RPMI 1640 medium. Plaques were cut into 2 mm^3^ fragments, mixed to ensure random distribution, and transferred into a 1-ml syringe for allocation into 48-well plates (0.1 ml tissue per well). Fragments were pre-incubated for 1 hour at 37°C in RPMI containing 10% fetal bovine serum under 5% CO_2_. Tissue was then stimulated with 20 ng/ml lipopolysaccharide for 1 hour, followed by treatment with the anti-FcRn monoclonal antibody rozanolixizumab or an isotype control. Incubations proceeded for 24 hours at 37°C. Supernatants were collected for MMP-9 and TNF quantification using the previously described ELISA assays. Residual plaque tissue was processed for protein extraction as outlined for frozen plaques, and total protein content was measured using the Bradford assay (Bio-Rad).

### Statistical Analysis

Statistical methods, sample sizes, and summary metrics are reported in the corresponding figures and legends. Data normality was evaluated using the Shapiro-Wilk test, and outliers were identified with the robust regression and outlier removal (ROUT) method (Q = 1%). All analyses were performed in GraphPad Prism 9.5.1, and statistical significance was defined as *p* < .05.

## Results

### Immunoglobulin Landscape Across Atherosclerotic Plaque Regions

Deposits of IgG, IgA, IgM, and IgE were detected by immunostaining in carotid plaques from 30 endarterectomy patients (Fig. 1A, S1A). Staining was predominantly extracellular but also present on cellular structures. IgG, IgA, and IgM were broadly distributed, whereas IgE was least abundant and localized mainly to scattered IgE^+^ cells (likely mast cells or macrophages) and deeper plaque regions adjacent to the tunica media. Among the isotypes, IgM showed the largest stained area, possibly reflecting recognition of a wider antigen spectrum or assay-related sensitivity.

**Figure 1.**
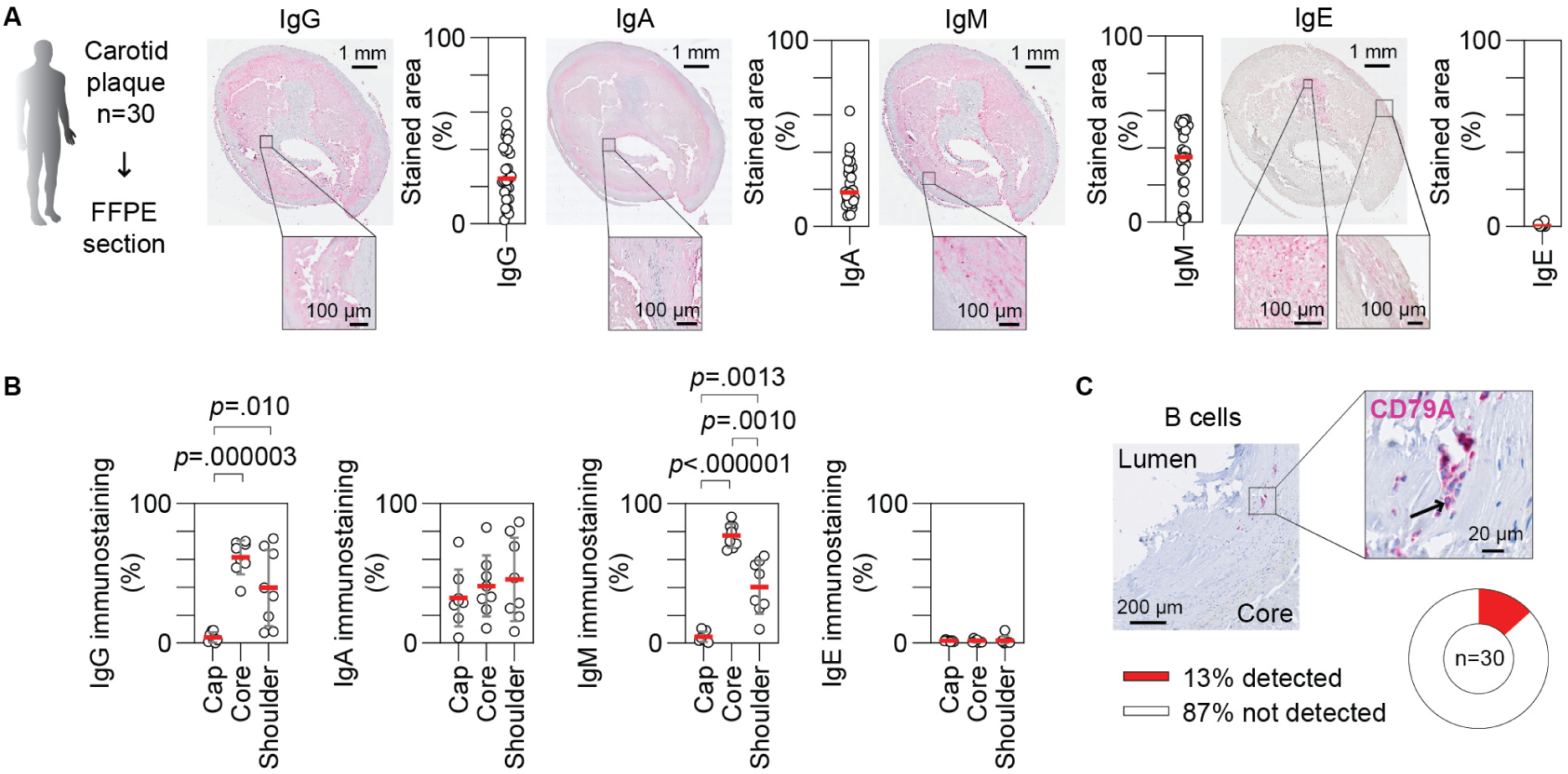
Spatial distribution of immunoglobulins in human atherosclerotic plaques. (**A**) Representative formalin-fixed paraffin-embedded (FFPE) carotid plaque sections stained for IgA, IgE, IgM, and IgG, with quantification of total antibody-positive area per plaque. (**B**) Regional analysis of antibody staining in core, shoulder, and fibrous cap compartments (repeated-measures one-way ANOVA with Tukey’s post hoc test; n = 8 plaques). (**C**) Representative CD79A staining highlighting intraplaque B cells, with frequency of B-cell detection across samples. Data are presented as mean ± SD and scale bars as indicated.

In plaques with preserved morphology, regional quantification revealed that IgA was evenly distributed, while IgG and IgM displayed stronger and more regionally distinct staining patterns (Fig. 1B, S1A-B). Although their patterns overlapped, IgM was more uniformly enriched in the necrotic core than in the inflammatory shoulder. To determine whether this reflected lipid peroxidation or necrotic material, we used LR04, an IgM antibody recognizing malondialdehyde-modified epitopes common to oxLDL and dead cells [35]. LR04 staining did not correlate with total IgM (Fig. S1C), indicating that IgM deposition was not solely driven by these processes. Notably, both IgM and IgG were significantly reduced in the fibrous cap relative to other plaque regions, suggesting antigen-poor zones or restricted immunoglobulin penetration. No sex differences in IgM or IgG deposition were observed (Fig. S1D).

To determine whether local B cells contributed to plaque antibody deposition, we performed B-cell immunostaining (Fig. 1C). B cells were rare, detected in only four plaques with 2-38 cells, indicating that most plaque immunoglobulins are unlikely to be locally produced. CD79A was used to identify B cells and plasma cells, and in the few positive plaques, B cells were located between the fibrous cap and necrotic core. Interestingly, B cells were more frequently detected in female patients (Fig. S1E), although overall antibody deposition did not differ by sex (Fig. S1D), arguing against a major role for local B-cell production in shaping the observed antibody patterns.

Antibodies primarily accumulated in deeper plaque regions rather than in the luminal cap, suggesting entry via intraplaque microvessels. In contrast, IgA showed uniform staining throughout the plaque, consistent with full tissue accessibility and possible accumulation according to antigen density. To explore mechanisms of immunoglobulin transport, we examined expression of known transport receptors. In single-cell ATAC-sequencing data from human atherosclerotic coronary arteries [36], FCGRT, PIGR, and FCER2 were predicted to be expressed by endothelial cells (Fig. S1F), although FCGRT was most strongly associated with a subset of plaque macrophages. Collectively, these findings suggest that transcytosis and/or transudation of systemic antibodies account for the bulk of plaque immunoglobulin content, with only limited contribution from local B-cell activity.

### Plaque and Plasma Immunoglobulin Profiles in Carotid Atherosclerosis

To further examine immunoglobulin distribution, we analyzed 72 paired plaque and plasma samples and normalized plaque antibody levels to total protein content (Fig. 2A). All four isotypes, IgG, IgA, IgM, and IgE, were detectable in plaques at concentrations that broadly reflected their relative abundance in plasma (Fig. 2B-C). Correlation analyses revealed moderate and comparable concordance between plaque and plasma IgA and IgM, consistent with their constitutive production and established reactivity toward oxLDL [37]. In contrast, plaque IgG did not correlate with plasma levels, likely reflecting the broader and stimulus-dependent specificity of circulating IgG. IgG, IgA, and IgM concentrations within plaques were positively interrelated, suggesting that certain lesions accumulate higher overall amounts of antigen or infiltrating plasma components (Fig. 2D).

**Figure 2.**
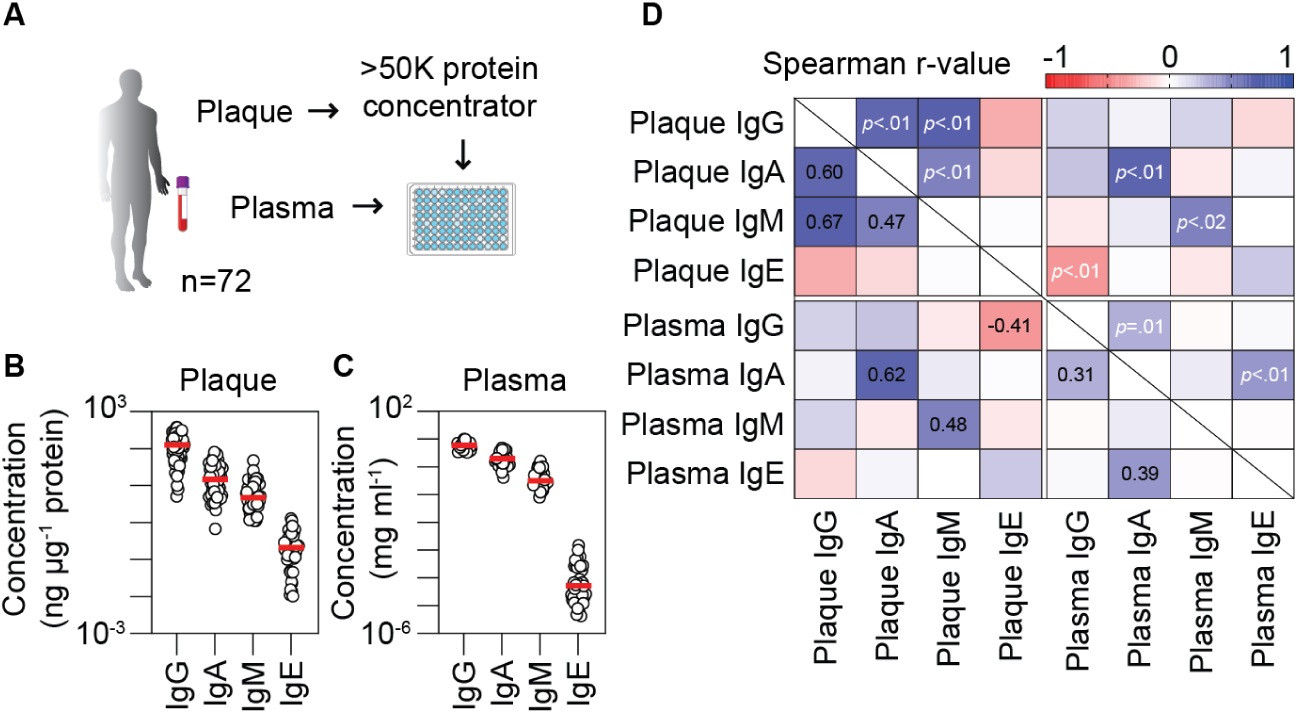
Antibody concentrations in human atherosclerotic plaques and corresponding plasma. (**A**) Schematic overview of the experimental workflow for plaque homogenization and immunoglobulin quantification. (**B**) Concentrations of immunoglobulin classes measured in homogenized carotid plaque extracts. (**C**) Matching immunoglobulin concentrations measured in patient plasma. (**D**) Heatmap illustrating correlations between plasma and plaque immunoglobulin levels; Spearman r-values are shown in the lower left matrix and *p*-values in the upper right. Data are presented with median levels indicated.

Females exhibited higher plasma IgM levels and a stronger correlation between plasma and plaque IgM than males (Fig. S2A-C), whereas no sex differences were observed for the other isotypes. Total immunoglobulin concentrations in plasma or plaques did not differ between asymptomatic patients and those with recent transient ischemic attack or stroke (Fig. S2D-E). Because IgE can vary with allergen exposure, we assessed seasonal patterns but found no variation in plasma or plaque IgE, or in the other isotypes, across the year (Fig. S3). Overall, plaque antibody levels were stable over time, largely reflecting systemic immunoglobulin availability rather than clinical status or environmental factors.

### Immunoglobulin Deposition Tracks with Structural and Inflammatory Markers of Plaque Vulnerability

To assess the pathological significance of plaque immunoglobulins, we performed morphological characterization of the carotid stenosis histology cohort. Antibody staining was inversely associated with collagen content, smooth muscle cell presence, and fibrous cap thickness (Fig. 3A-B, S4A-E). This pattern was most pronounced for IgG and IgM, although both were generally sparse in the cap where α-smooth muscle actin-positive cells reside. Antibody staining correlated positively with T-cell abundance but not with B-cell counts, further supporting that most plaque immunoglobulins arise from transudation or transcytosis rather than local production.

**Figure 3.**
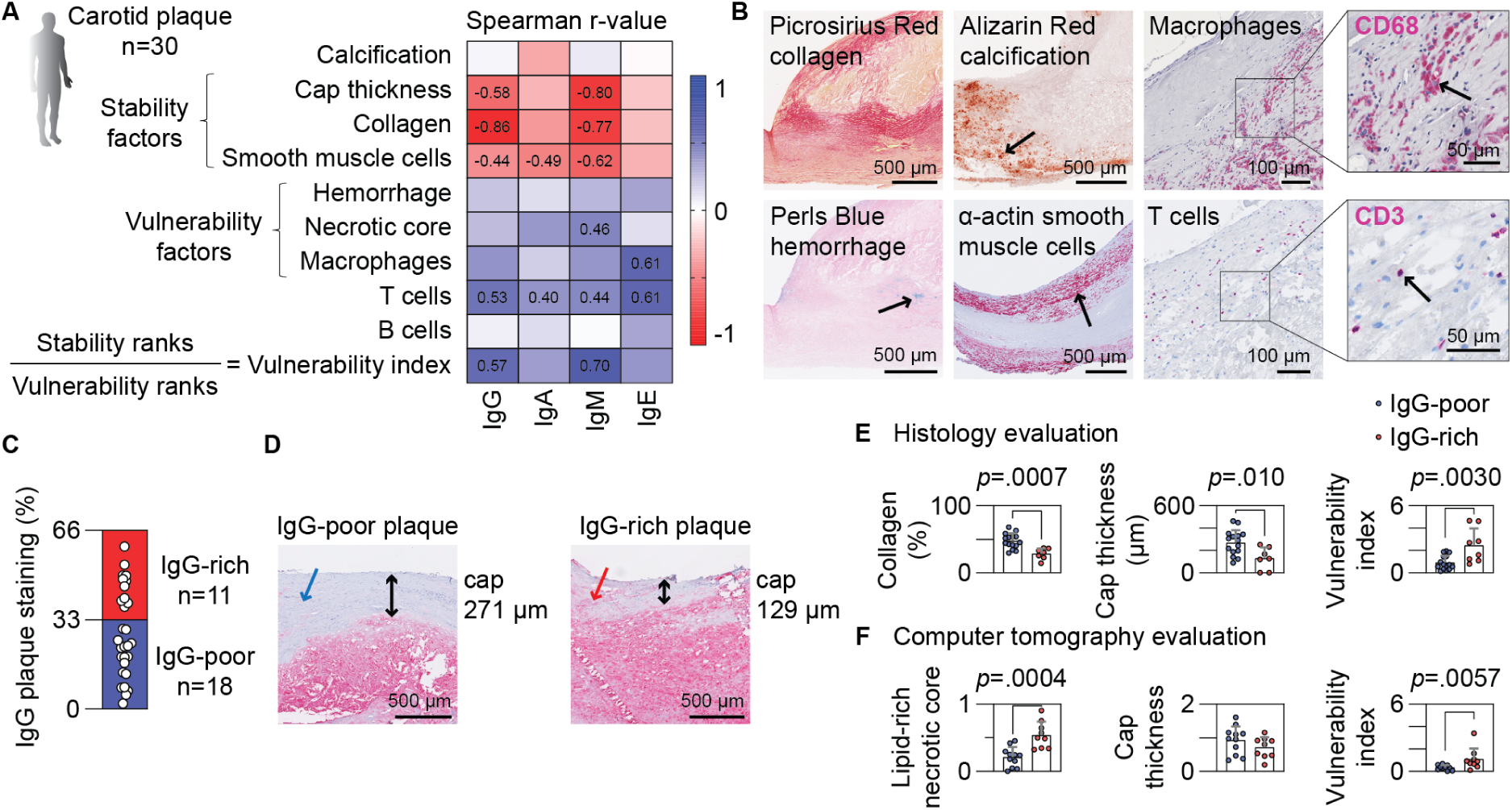
Local IgG deposition associates with markers of atherosclerotic plaque vulnerability. (**A**) Heatmap showing correlations between IgG staining and morphological features of plaque vulnerability; Spearman r-values are displayed for significant correlations. (**B**) Representative micrographs illustrating the assessed morphological features. (**C**) Stratification of plaques into IgG-poor (<33% IgG⁺ area) and IgG-rich (>33% IgG⁺ area) groups. (**D**) Representative cap-region micrographs from IgG-poor and IgG-rich plaques; colored arrows indicate cap structure, and black arrows denote measured cap thickness. (**E**) Quantification of collagen content, cap thickness, and vulnerability index in IgG-rich versus IgG-poor plaques (unpaired *t-*test). (**F**) Computer tomography-based quantification of lipid-rich necrotic core, minimal cap thickness, and vulnerability index in IgG-rich versus IgG-poor plaques (Mann-Whitney test). Data are presented as mean ± SD and scale bars as indicated.

The overall vulnerability index was associated with IgG and IgM, but not with IgA or IgE (Fig. 3A). Plaque IgG and IgM were strongly correlated with each other (Spearman r = 0.86; Fig. S4F), making independent associations difficult to disentangle. Notably, plaque immunoglobulin deposition did not correlate with traditional cardiovascular risk factors, including age, sex, body mass index, diabetes, hypertension, hypercholesterolemia, or plasma high-sensitivity C-reactive protein (Fig. S4G), suggesting that antibody burden reflects a biologically distinct vulnerability feature not captured by standard risk metrics.

To illustrate the heterogeneity, plaques were stratified by IgG content. IgG-poor plaques (≤1/3 IgG-positive area) displayed thicker fibrous caps with greater collagen deposition (Fig. 3C-E). In contrast, IgG-rich plaques exhibited thinner caps, higher vulnerability indices, and increased NLRP3⁺ cell density (Fig. S4H). Necrotic-core area did not differ by IgG burden; instead, IgG-rich plaques frequently showed antibody infiltration into fibrotic and shoulder regions (Fig. 3D), placing IgG in direct proximity to immune and stromal cells within inflamed microenvironments. Complementary computed tomography analyses indicated greater vulnerability in IgG-rich plaques, driven primarily by larger lipid-rich necrotic cores rather than differences in cap thickness or intraplaque hemorrhage (Fig. 3F, S3I).

### ApoB-Specific IgG Accumulation and Immune Complex Dynamics in Plaques

Plaque apoB levels positively associated with immunoglobulin deposition, most notably IgG (Fig. 4A), suggesting that a portion of lesional IgG reflects binding to apoB-containing lipoproteins. Consistent with this, the fraction of anti-apoB IgG was higher in plaque than in plasma (Fig. 4B), whereas anti-apoB IgM was reduced (Fig. S5A). Symptomatic patients exhibited elevated plaque anti-apoB IgG, an effect not observed in blood (Fig. 4C, S5B-C), pointing to local enrichment of this specificity (Fig. 4D). In line with increased local immune complex turnover, plaques from symptomatic patients contained less apoB and fewer apoB-IgG immune complexes (Fig. 4E-F), and plaque immune complexes did not correlate with their plasma counterparts (Fig. S5D-F). Among interrelationships, the strongest was an inverse correlation between plaque apoB and anti-apoB IgG (Fig. 4G), consistent with enhanced macrophage-mediated immune complex uptake within lesions.

**Figure 4.**
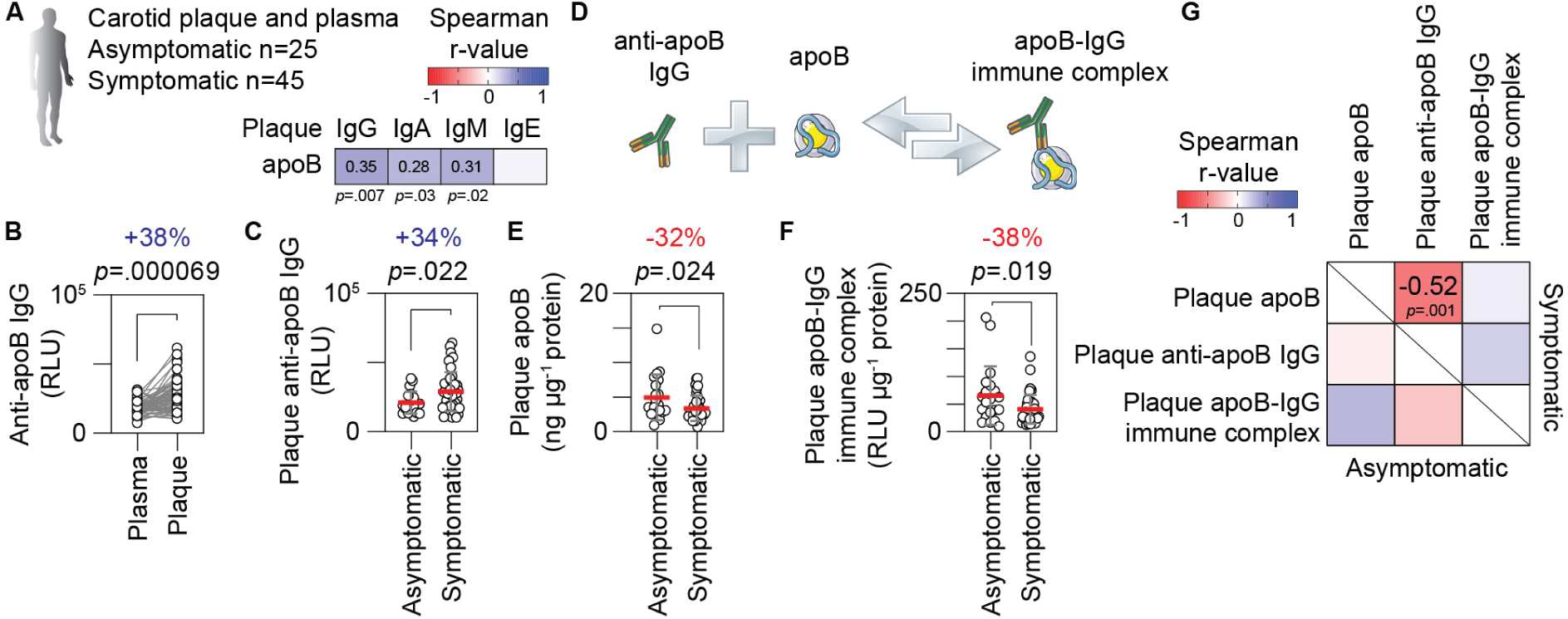
Dynamic accumulation of apoB-specific IgG in human atherosclerotic plaques. (**A**) Associations between immunoglobulin concentrations and apoB levels in carotid plaque protein extracts. (**B**) Anti-apoB IgG levels measured in paired plasma and plaque samples normalized to 1 µg/ml total IgG (paired *t*-test). (**C**) Plaque anti-apoB IgG levels in asymptomatic versus symptomatic patients (unpaired *t*-test). (**D**) Schematic illustration of apoB, anti-apoB IgG, and apoB-IgG immune complexes. (**E**) Plaque apoB concentrations (unpaired *t*-test). (**F**) Levels of apoB-IgG immune complexes in plaques (unpaired *t-*test). (**G**) Matrix illustrating relationships among these parameters in asymptomatic (lower left) and symptomatic (upper right) patients. RLU, relative light units. Data are presented as mean ± SD.

In plaques from symptomatic patients, apoB-containing lipoproteins were relatively depleted of IgG1 compared with IgG2-4 (Fig. S5G-H). Plaque apoB, anti-apoB IgG, and apoB-IgG immune complexes were not associated with traditional risk factors (body mass index, diabetes, hypertension, plasma lipids, high-sensitivity C-reactive protein; Fig. S5I-J). However, plaque apoB and apoB-IgG complexes declined with age, while plaque anti-apoB IgG increased, patterns not seen in plasma, implicating age-linked local processes. Plaque apoB-IgG complexes were also associated with male sex, consistent with prior plasma observations [38].

### Age-associated Expansion of FcRn^+^Macrophages in Human Atherosclerosis

FCGRT mRNA, which encodes the IgG recycling receptor FcRn, was markedly upregulated in carotid plaques compared with non-diseased arterial tissue (Fig. 5A). Immunofluorescence localized FcRn protein to CD163^+^ macrophages in plaque shoulder regions (Fig. 5B). Integrating single-cell ATAC and RNA sequencing from human atherosclerotic arteries identified a macrophage subset with the highest FCGRT signal (Fig. S6A), a pattern recapitulated in mouse atherosclerotic tissue, where Fcgrt was abundant in Cd163^+^ macrophages with resident-like features (Fig. S6B-C). To link IgG handling with vulnerability programs [39], we examined matrix-remodeling enzymes in the same cells: Mmp9 showed a congruent expression pattern (Fig. S6D), suggesting a macrophage phenotype capable of both IgG recycling and matrix remodeling.

**Figure 5.**
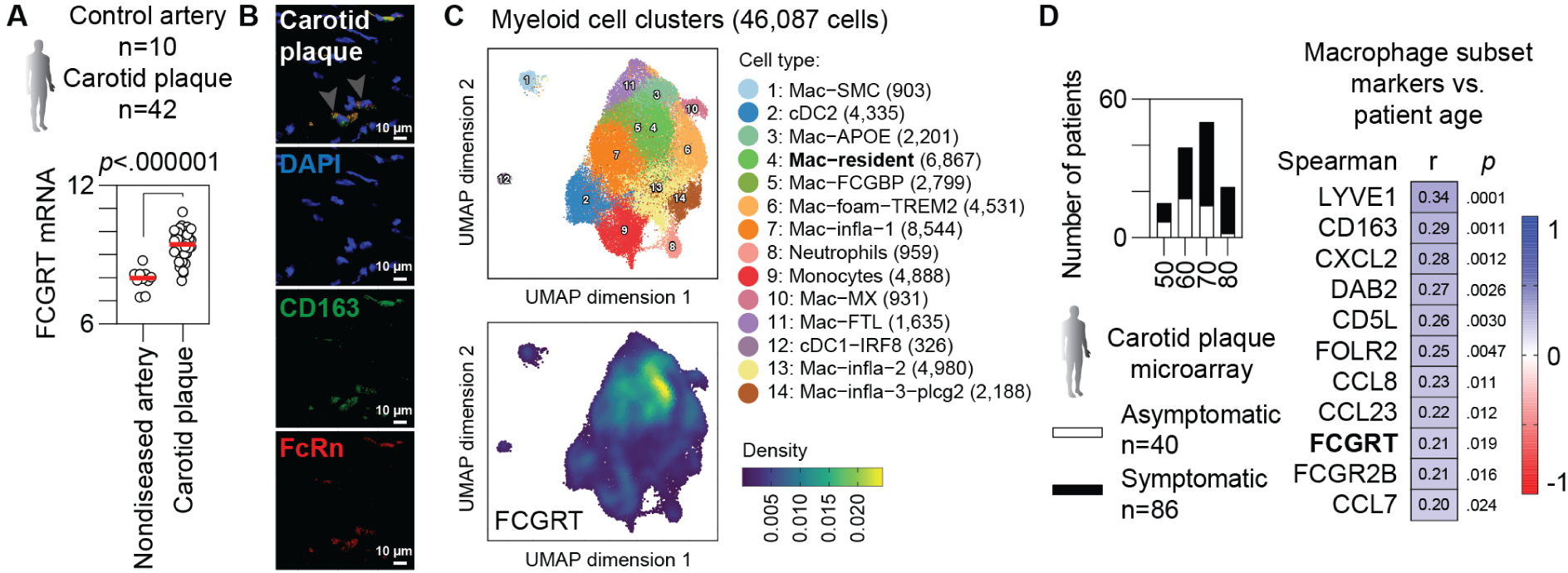
Expression of the IgG recycling receptor FcRn in plaque macrophages. (**A**) FCGRT expression in carotid plaques compared with non-diseased iliac arteries (unpaired *t*-test). Data are presented with median levels indicated. (**B**) Immunofluorescence staining of FcRn and CD163⁺ macrophages in the shoulder region of a carotid plaque. (**C**) Integrated single-cell RNA-seq analysis of myeloid populations in human atherosclerosis with a feature plot showing FCGRT expression (n = 26). (**D**) Age distribution in the carotid stenosis cohort and Spearman correlations between macrophage subset markers and patient age. Mac, macrophage; cDC, conventional dendritic cell; infla, inflammatory.

Integration of human single-cell datasets provided additional support for FCGRT^+^ macrophages in advanced lesions (Fig. 5C). Reclustering myeloid cells into 14 subsets revealed that FCGRT expression was concentrated in a resident-like macrophage cluster. In our carotid endarterectomy cohort, FCGRT mRNA showed a modest positive association with age, along with ten markers of resident-like macrophage states (LYVE1, CD163, CXCL2, DAB2, CD5L, FOLR2, CCL8, CCL23, FCGR2B, and CCL7; Fig. 5D, S6E-F). These markers were mutually correlated (Fig. S6G), suggesting that this FCGRT^+^ macrophage population increases with age, notable given that symptomatic patients were, on average, 6 years older than the asymptomatic patients.

Reanalysis of the dataset from Lee et al. [33] further showed higher FCGRT expression in laser-microdissected macrophage-rich regions of ruptured human plaques compared with stable lesions (Fig. S6H). Taken together, these results point toward age-associated expansion of FcRn-expressing macrophages in atherosclerosis and are consistent with a potential role for IgG-handling pathways in plaque progression.

### FcRn-Mediated Recycling of Anti-apoB IgG in Macrophages

To examine whether FcRn mediates recycling of anti-apoB IgG, we performed in vitro assays using RAW 264.7 macrophages (Fig. 6A). IgG enriched for anti-human apoB reactivity, and cross-reactive to oxLDL, was isolated from *APOB100*-transgenic mice that had received apoB-reactive T cells (Fig. S7A). Macrophages were cultured on plates coated with immune complexes formed from these IgGs, and Fcgrt silencing efficiently reduced Fcgrt mRNA without inducing cytotoxicity (Fig. 6B, S7A). Knockdown of *Fcgrt* led to lower IgG levels in the supernatant, consistent with reduced antibody recycling, and was accompanied by decreased *Tnf*, *Il1b*, and *Mmp9* expression and reduced TNF secretion (Fig. 6B, S7B). *Fcgrt* silencing also produced a significant reduction in LDL uptake (Fig. 6B, S7C). Collectively, these results indicate that FcRn contributes to the recycling of anti-apoB immune complexes and is linked to macrophage inflammatory activation and lipoprotein handling in vitro.

**Figure 6.**
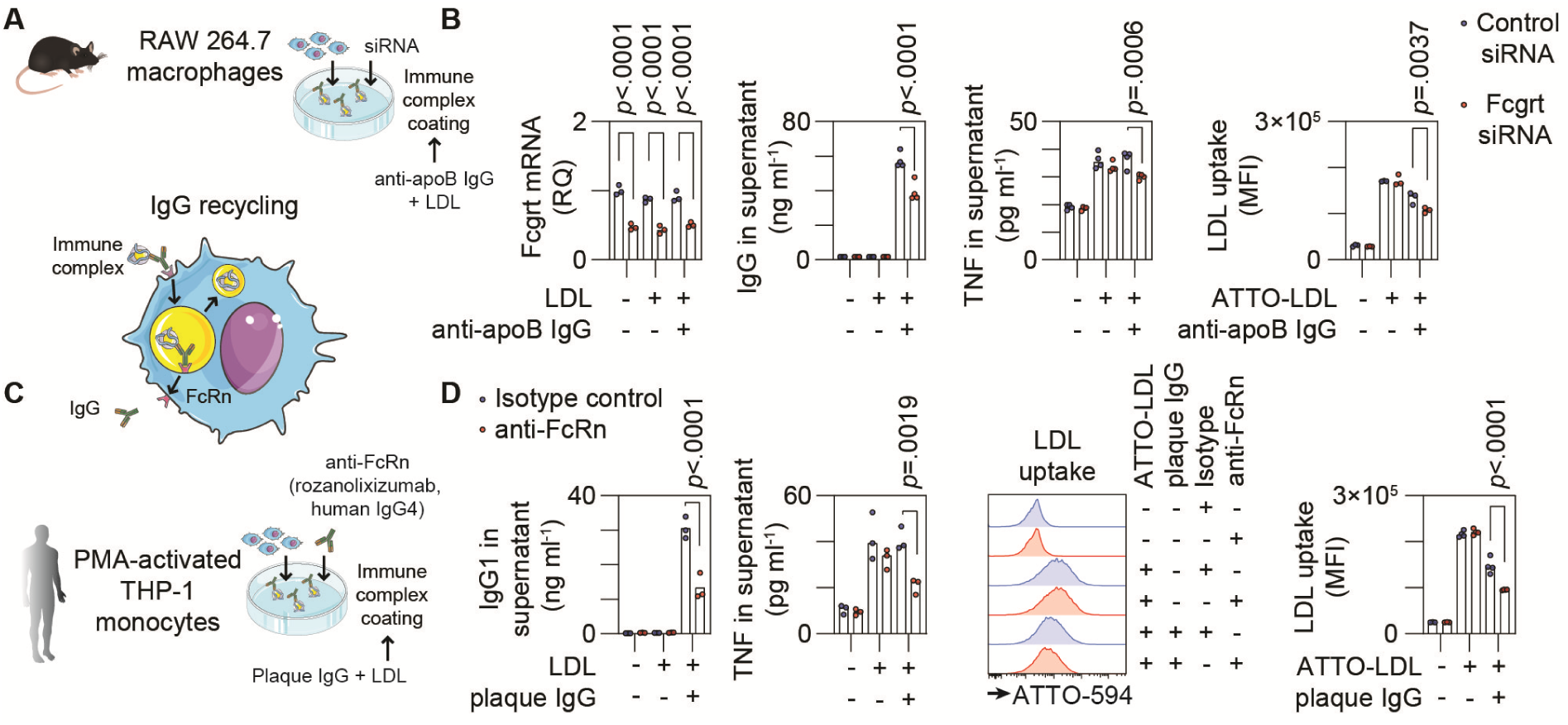
FcRn blockade or knockdown suppresses IgG recycling and inflammatory activation in macrophages. (**A**) Formation of LDL-IgG immune complexes in vitro using mouse IgG reactive against LDL. (**B**) Effects of Fcgrt knockdown in RAW 264.7 macrophages on IgG recycling and cytokine secretion, along with uptake of fluorescently labeled LDL (one-way ANOVA with Holm-Šídák correction). (**C**) Schematic of the THP-1 macrophage system used to assess the impact of rozanolixizumab on FcRn-dependent handling of immune complexes formed with plaque-derived IgG. (**D**) FcRn blockade reduces IgG1 recycling, TNF secretion, and alters uptake of fluorescently labeled LDL in THP-1 macrophages (one-way ANOVA with Holm-Šídák correction). MFI, mean fluorescence intensity; RQ, relative quantification. Data are presented with mean levels indicated.

We next assessed FcRn-dependent immune complex turnover in human macrophages, taking advantage of the availability of clinical-grade FcRn-blocking antibodies (Fig. 6C). IgG isolated from human plaques, showing cross-reactivity to apoB and oxLDL, was used to generate immune complex-coated plates, onto which THP-1-derived macrophages were added (Fig. S7D). The FcRn-blocking antibody rozanoliximab markedly inhibited recycling of IgG1, whereas IgG2 and IgG3 were less affected (Fig. 6D, S7D). FcRn blockade also reduced *TNF*, *IL1B*, and *MMP9* expression and lowered TNF secretion (Fig. 6D, S7E). In addition, rozanoliximab reduced LDL uptake in THP-1 macrophages without affecting cell viability (Fig. 6D, S7F). Together with the murine macrophage experiments, these findings support a role for FcRn in modulating IgG immune complex handling, inflammatory activation, and lipoprotein uptake in human macrophages.

### FcRn-linked MMP-9 Programs and Plaque Vulnerability

Single-cell analyses indicated Mmp9 expression within FcRn^+^ macrophages (Fig. S6D–E). In vitro, *Fcgrt* knockdown reduced Mmp9 mRNA and protein in immune complex-stimulated RAW 264.7 macrophages (Fig. 7A, S8A), and rozanoliximab similarly decreased MMP-9 expression and secretion in plaque antibody-stimulated THP-1 macrophages (Fig. 7B, S8B-C), consistent with transcriptional regulation. In plaques, MMP-9 protein positively correlated with anti-apoB IgG, with a stronger relationship in symptomatic cases (Fig. 7C). FCGRT mRNA correlated inversely with stability markers and positively with vulnerability factors (Fig. 7D), including a strong association with MMP9 (Spearman r = 0.72), and with TNF and IL1B mRNA levels, aligning with the reductions observed following FcRn blockade in vitro. Finally, in an ex vivo plaque culture, rozanoliximab decreased IgG recycling and reduced secretion of TNF and MMP-9 after lipopolysaccharide stimulation (Fig. 7E, S8D). Together, these data are consistent with a model in which FcRn-dependent IgG handling in macrophages is linked to MMP-9-associated vulnerability programs.

**Figure 7.**
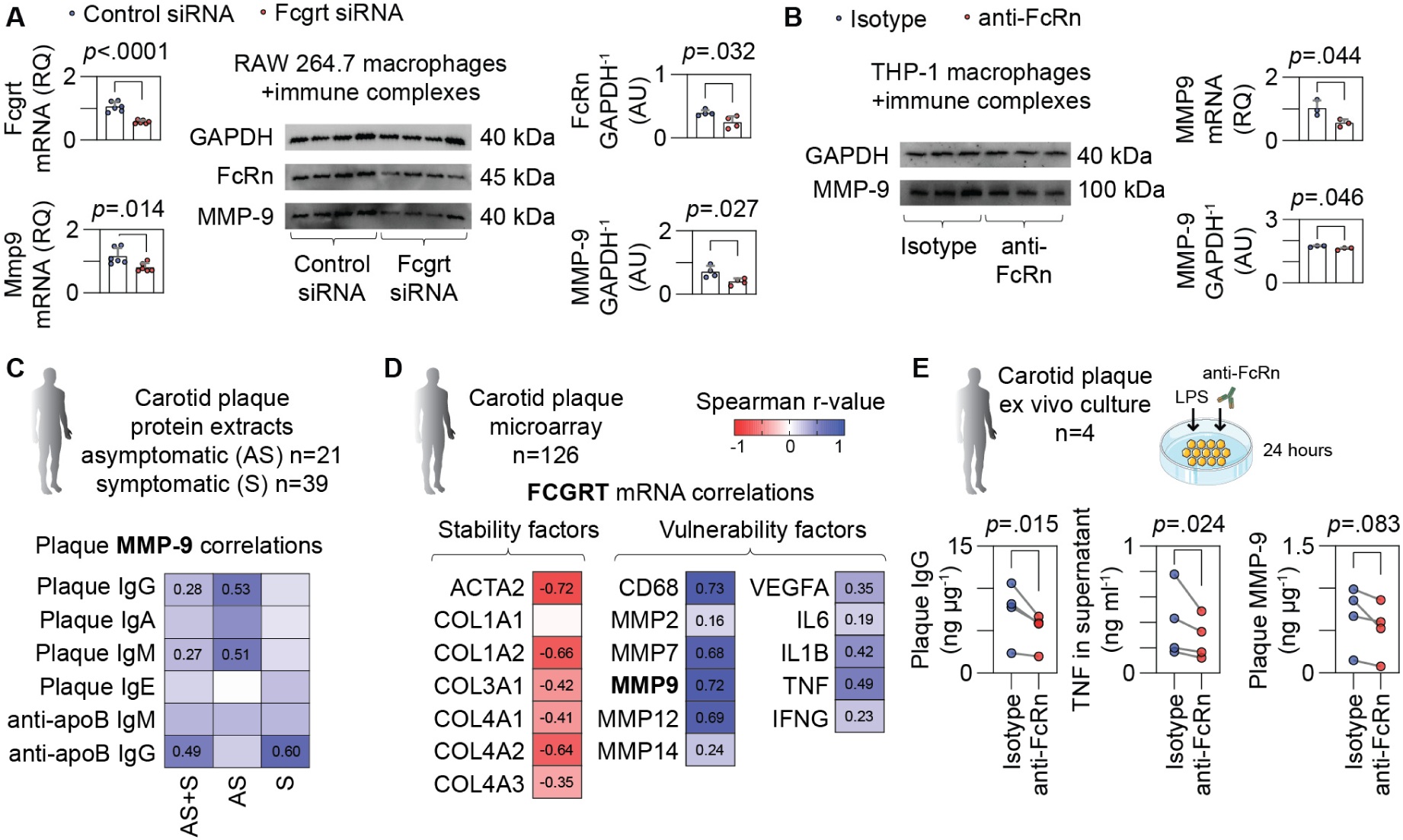
FcRn-dependent IgG recycling promotes MMP-9 production in macrophages and human plaques. (**B**) Silencing Fcgrt reduces MMP-9 production in RAW 264.7 macrophages stimulated with LDL immune complexes (unpaired *t*-test). (**C**) FcRn blockade with rozanolixizumab lowers MMP-9 production in LDL immune complex-stimulated THP-1 macrophages (unpaired *t*-test). (**D**) MMP-9 levels positively correlate with antibody concentrations in carotid plaques; Spearman r-values are shown for significant associations. (**E**) FCGRT transcript levels correlate with markers of plaque stability and vulnerability in carotid plaque tissue. (**F**) Ex vivo stimulation of human carotid plaques for 24 hours: IgG and MMP-9 were measured in plaque extracts, and TNF was quantified in supernatants (ratio paired *t*-test). AU, arbitrary units; RQ, relative quantification; AS, asymptomatic; S, symptomatic. Data are presented as mean ± SD.

## Discussion

We identify abundant immunoglobulins within atherosclerotic plaques as a correlate of plaque vulnerability and propose a mechanism in which FcRn-mediated antibody recycling in plaque macrophages increases with age and actively amplifies local inflammation and matrix degradation. This model explains (i) regional enrichment of IgG in macrophage-dense shoulder regions, (ii) elevated anti-apoB IgG in plaques, particularly in symptomatic patients, concurrent with reduced apoB immune complexes, and (iii) enhanced production of MMP-9 and pro-inflammatory cytokines by macrophages within the plaque. Together, these findings highlight FcRn as a therapeutic target within the inflammatory component of human atherosclerosis. Notably, FcRn blockade, already approved for generalized myasthenia gravis and in trials for other autoimmune diseases [40], reduced MMP-9 in our models, supporting a causal link between IgG recycling and cap-weakening biology and outlining a plausible path to translation.

We observed no sex-related differences in plaque antibody levels, despite higher circulating IgM in females and the well-established female bias in many autoimmune and inflammatory disorders [41]. This aligns with modern cardiovascular mortality statistics, which show comparable atherosclerosis related deaths between women and men [42]. Although female plaques contained more B cells, these lesions did not show enhanced antibody-driven inflammation. This finding indicates that antibody dependent mechanisms remain viable therapeutic targets in both sexes and suggests that FcRn mediated processes operate independently of the systemic immunologic dimorphism observed in other chronic inflammatory conditions.

The precise routes through which antibodies enter atherosclerotic plaques remain incompletely understood. Our findings support a model in which active endothelial transport, potentially mediated by specific receptors, complements passive entry associated with endothelial disruption, erosion, or mesenchymal transformation [43]. Intraplaque hemorrhage represents another theoretical route, yet plaques with hemorrhage did not show higher antibody content in our analyses. Instead, our data point toward FcRn-mediated IgG recycling as a major contributor to antibody accumulation within plaques, a process likely amplified in macrophage-dense regions and further accentuated with aging, which we found to be associated with expansion of the FcRn^+^ macrophage population.

FcRn plays a central role in maintaining IgG levels in tissues by binding the Fc domain in acidic endosomes, rescuing IgG from lysosomal degradation, and releasing it at neutral pH [44]. In atherosclerotic plaques, this pathway provides a mechanism for IgG to be transported across endothelial barriers and delivered to cognate antigens. Through its pH-dependent binding cycle, FcRn diverts IgG-containing vesicles away from degradation and returns them to the cell surface or across the cell via transcytosis.

FcRn displays subclass-specific affinity, with IgG1 exhibiting the highest recycling efficiency [44]. This hierarchy aligns with our finding of abundant apoB-IgG1 immune complexes within carotid plaques and the pronounced FcRn-dependent recycling of plaque-derived IgG1 in vitro. Plasma levels did not mirror those in plaques, indicating that systemic measurements and intraplaque immune complex burdens are uncoupled, consistent with rapid reticuloendothelial clearance of circulating complexes. Immune complex handling therefore appears to be locally regulated: a subset of plaque macrophages expressed FcRn, identifying them as the primary cells mediating IgG recycling within lesions. In contrast, foamy macrophages lacked FcRn expression, indicating minimal involvement in antibody recycling. Nonetheless, FcRn contributed to LDL uptake in macrophages in vitro, indicating that it may influence both immune and metabolic macrophage functions within the plaque.

Prior work has shown that regional IgG accumulation in plaque tissue can serve as a marker of inflammation and may enable advanced molecular imaging strategies [45]. Such an approach may gain therapeutic relevance if IgG-enriched plaques can be selectively targeted with FcRn-blocking agents, enabling a precision-medicine strategy focused on lesions with excessive antibody recycling. In this context, our classification of plaques into IgG-rich and IgG-poor states could complement existing methods used to distinguish inflammatory “hot” from quiescent “cold” plaques, paralleling frameworks used in tumor immunology to identify likely responders to immunomodulatory therapies.

FcRn blockade may interrupt a self-reinforcing cycle of antibody recycling and inflammation. In human plaques, FcRn expression was associated with increased levels of pro-inflammatory mediators, including TNF and IL-1β, and co-regulated with MMP-9, a key effector of fibrous-cap degradation. Elevated NLRP3 inflammasome expression further supports ongoing IL-1β activation within these lesions. In addition, experimental data presented in abstract form indicate that macrophage Fcgrt expression accelerates atherosclerosis progression and vascular inflammation in *Ldlr*^-/-^ mice, further implicating FcRn in disease pathogenesis [46].

Macrophages shape every stage of atherosclerosis through roles in lipid handling, matrix remodeling, endocytosis, and efferocytosis, and shifts in their functional states can accelerate disease progression [47, 48]. With aging, plaques exhibit an increased representation of inflammatory macrophage subsets [49], aligning with our observation that FcRn^+^ macrophages are positioned to mediate pathogenic IgG recycling in advanced lesions.

The link between enhanced IgG recycling and symptomatic disease raises important questions about how this mechanism contributes to plaque vulnerability. IgG is the dominant immunoglobulin within plaques, and prior studies have shown that IgG can exert pro-inflammatory and pro-atherogenic effects [50, 51]. Our findings suggest that accelerated antibody recycling by FcRn^+^ macrophages may amplify these responses and promote vulnerability-related pathways, including MMP-9-associated matrix degradation. The magnitude of this contribution warrants further investigation, but these observations highlight FcRn^+^ macrophages as a mechanistically relevant population driving IgG-dependent inflammation in high-risk plaques.

Although IgG and IgM have been well characterized in atherosclerosis, the roles of IgA and IgE within plaques remain insufficiently explored [52]. Plaque microenvironments also exhibit pH heterogeneity, which may influence FcRn-IgG interactions and deserves further evaluation [53]. While our ex vivo model captures aspects of the plaque milieu, in vitro systems cannot fully recapitulate the structural and immunologic complexity of human lesions. Longitudinal studies will be required to determine whether FcRn activity contributes to plaque progression, and interventional studies will ultimately be needed to define its causal impact on disease.

In conclusion, our findings identify FcRn-dependent handling of apoB-immune complexes by FcRn^+^ macrophages as a mechanistic driver of plaque vulnerability, positioning FcRn as a potential therapeutic target in atherosclerosis.

## Author contributions

SL and AG compiled and analyzed the data and wrote the manuscript. The authors declare no conflict of interest. We thank Linda Haglund for technical support.

## Funding

This work was supported by grants from the Swedish Heart-Lung Foundation (20210469 and 20230391), Swedish Research Council (2020-01789), Swedish Society of Medicine, Leducq Foundation (TNE-20CVD03), Åke Wibergs Stiftelse, Jeanssons Stiftelse, Magnus Bergvalls Stiftelse, Foundation for Age Research at Karolinska Institutet, Karolinska Institutet Research Foundation, Eva and Oscar Ahréns stiftelse, Stiftelsen Professor Nanna Svartz, Stiftelsen för Gamla Tjänarinnor, the Finnish Foundation for Cardiovascular Research, and Aarne Koskelo Foundation.

## Notes

### Competing Interest Statement

The authors have declared no competing interest.

