## Supplemental Figures and Tables for "Antibody recycling via FcRn drives atherosclerotic plaque vulnerability"

**Figure S1**. Validation of antibody staining, sex-related differences in plaque immunostaining, and endothelial expression of immunoglobulin transport receptors.

**Figure S2**. Sex- and symptom-related variation in antibody concentrations in plasma and plaques.

**Figure S3**. Seasonal variation in antibody concentrations in plasma and plaques.

**Figure S4**. Associations between plaque antibody staining, plaque morphology, and clinical characteristics.

**Figure S5**. Associations between apoB-specific antibodies, clinical symptoms, and patient characteristics.

**Figure S6**. Expression of the neonatal Fc receptor in resident plaque macrophages.

**Figure S7**. FcRn blockade suppresses IgG recycling and inflammatory activation in macrophages.

**Figure S8**. MMP-9 production in macrophages and human atherosclerotic plaques.

*Supplemental Tables*

**Table S1**. Characteristics of patients with advanced carotid atherosclerosis included in the study.

**Table S2**. Antibodies used for immunohistochemistry, including clone, source, and working dilution.

**Table S3**. Antibodies used for immunoassays, including capture, detection, and isotype-specific reagents.

**Table S4**. Assay-on-demand probes and siRNA reagents used for real-time PCR and gene-silencing experiments.

### Supplemental Figures

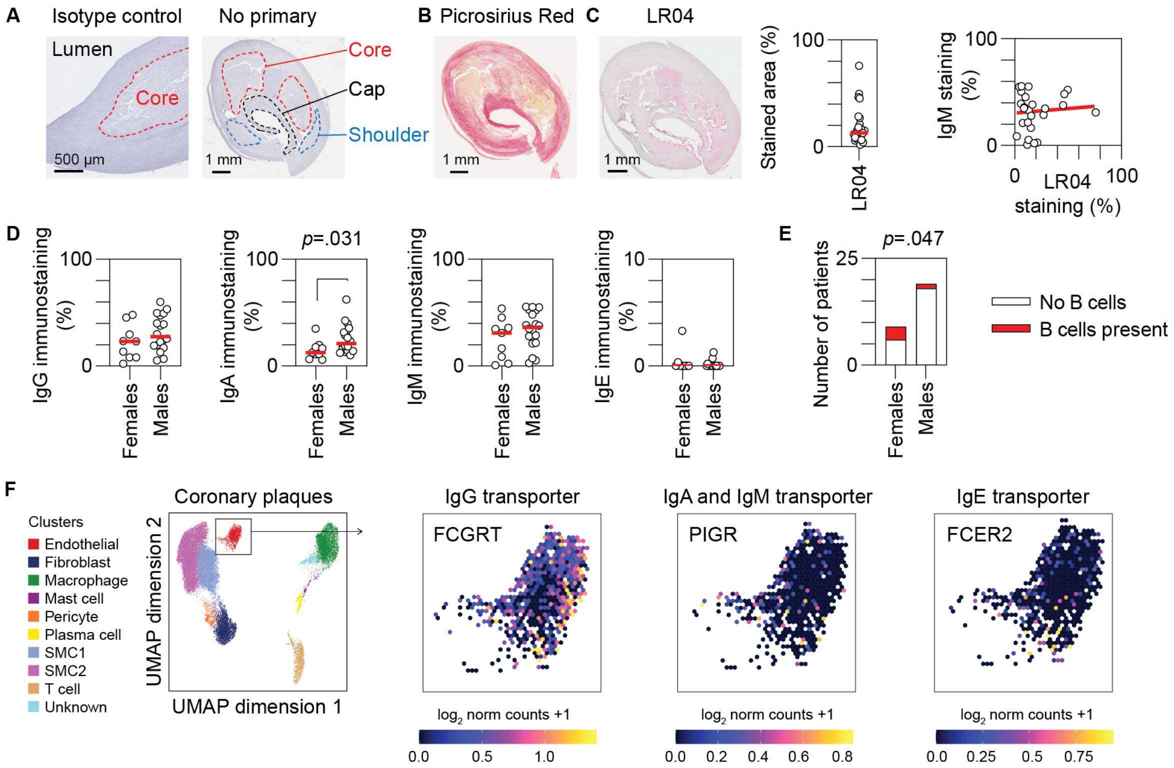


**Figure S1.** Validation of antibody staining, sex-related differences in plaque immunostaining, and endothelial expression of immunoglobulin transport receptors.
(**A**) Isotype control staining demonstrating specificity of antibody labeling and delineation of plaque regions in a section processed without primary antibody.
(**B**) Picrosirius Red staining used to define structural regions of the plaque.
(**C**) LR04 staining with quantification and linear regression analysis correlating LR04 signal with total IgM staining.
(**D**) Sex-related differences in plaque antibody distribution (unpaired *t*-test).
(**E**) Sex-related differences in plaque B-cell presence (Chi-square test).
(**F**) A single-cell assay for transposase-accessible chromatin sequencing dataset of human coronary atherosclerotic plaque endothelial cells accessed via the PlaqView portal, showing uniform manifold approximation and projection (UMAP) visualization of predicted gene scores for FCGRT, PIGR, and FCER2 across endothelial cells from 41 coronary artery disease patients (Turner et al., *Nature Genetics*, 2022).

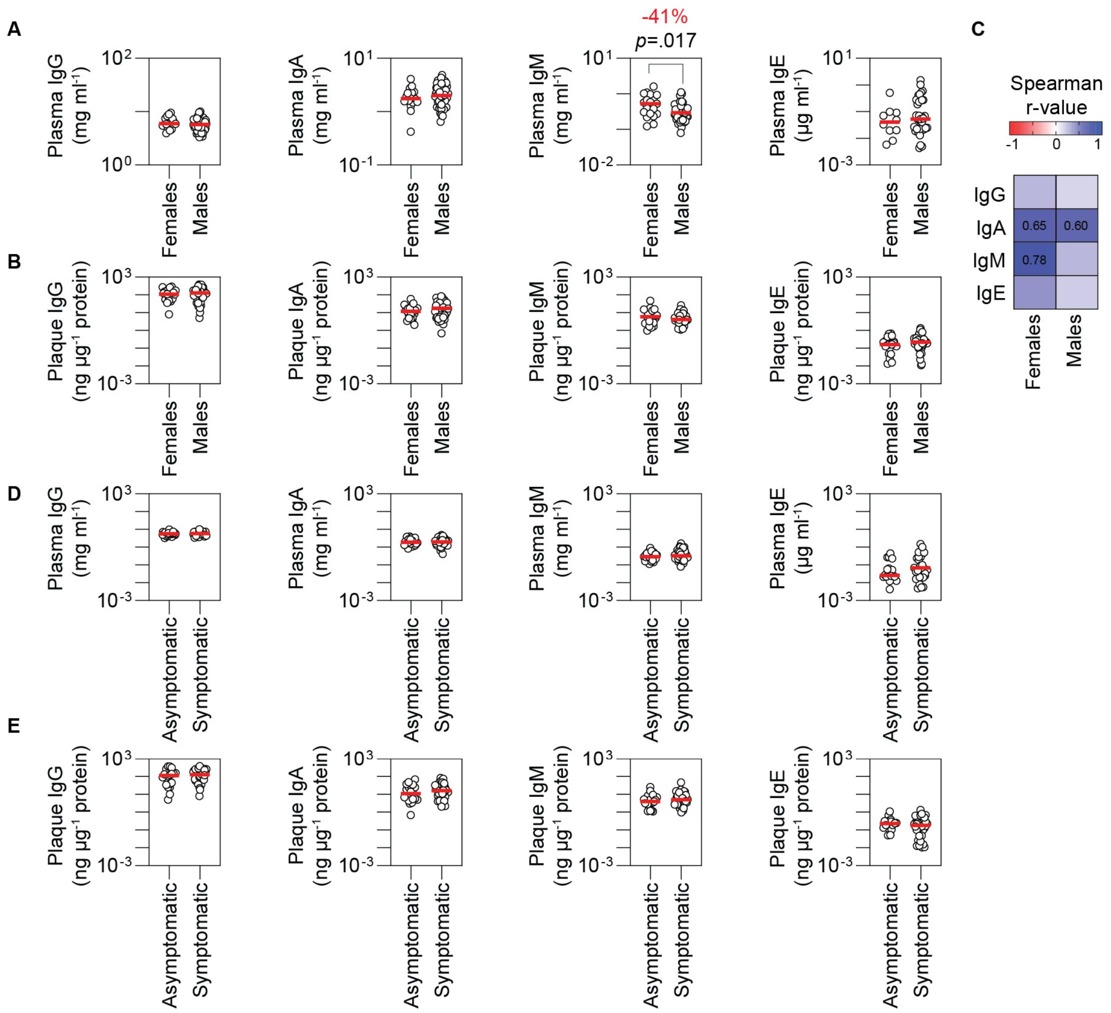


**Figure S2.** Sex- and symptom-related variation in antibody concentrations in plasma and plaques.
(**A**) Plasma antibody concentrations in females versus males (unpaired *t*-test).
(**B**) Antibody concentrations in carotid plaque extracts stratified by sex.
(**C**) Heatmap showing correlations between plasma and plaque antibody concentrations stratified by sex; Spearman r-values displayed for significant associations ($p < .05$).
(**D**) Plasma antibody concentrations in asymptomatic versus symptomatic patients.
(**E**) Corresponding antibody concentrations in carotid plaques from asymptomatic and symptomatic patients.

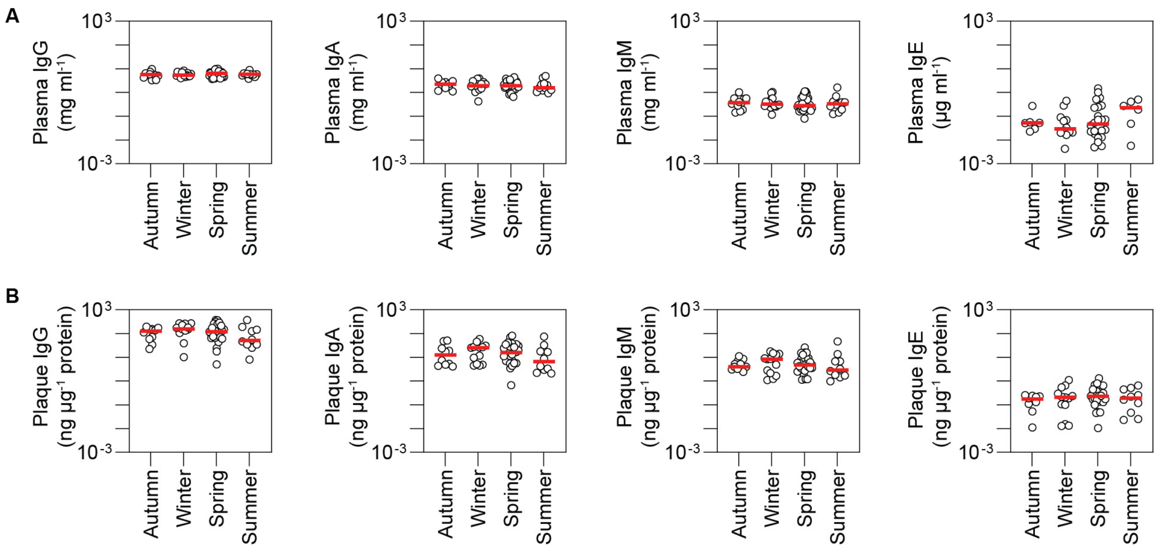


**Figure S3.** Seasonal variation in antibody concentrations in plasma and plaques.
(**A**) Seasonal variation in plasma concentrations of IgG, IgA, IgM, and IgE.
(**B**) Seasonal variation in corresponding antibody concentrations within carotid plaque extracts.

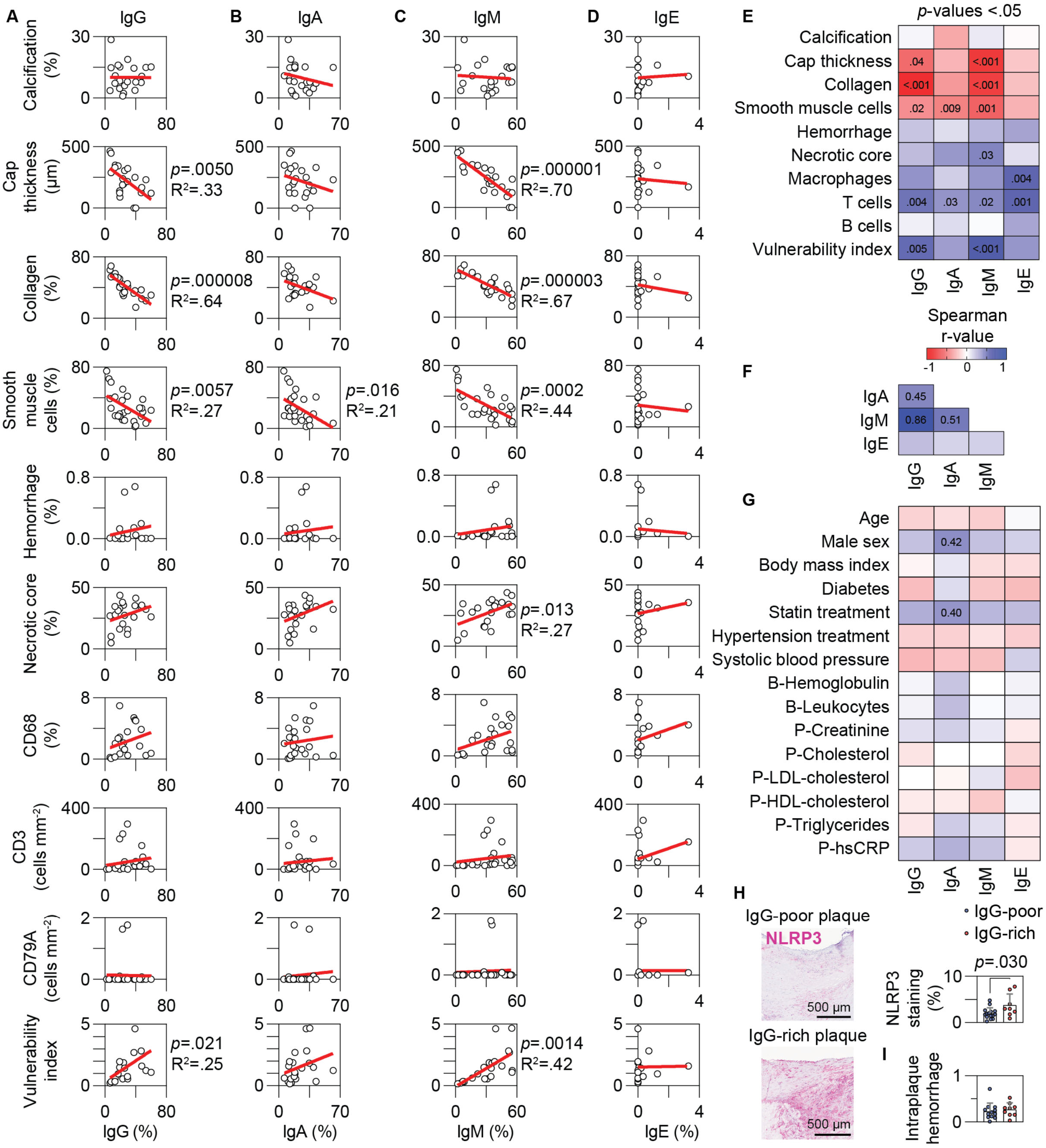


**Figure S4.** Associations between plaque antibody staining, plaque morphology, and clinical characteristics.

(**A-D**) Linear regression analyses illustrating relationships between morphological plaque features and plaque distribution for IgG (**A**), IgA (**B**), IgM (**C**), and IgE (**D**).

(**E**) Significant *p*-values corresponding to the correlations shown in the heatmap in Figure 3A between antibody staining and morphological plaque features.

(**F**) Heatmap showing inter-correlations among antibody stainings; Spearman r-values are shown for significant associations ($p < .05$).

(**G**) Heatmap showing correlations between antibody staining and clinical parameters; Spearman r-values displayed for significant associations ($p < .05$).

(**H**) Quantification of NLRP3 staining in IgG-poor and IgG-rich plaques, with representative micrographs (unpaired *t*-test).

(**I**) Computer tomography-based quantification of intraplaque hemorrhage in IgG-rich versus IgG-poor plaques.

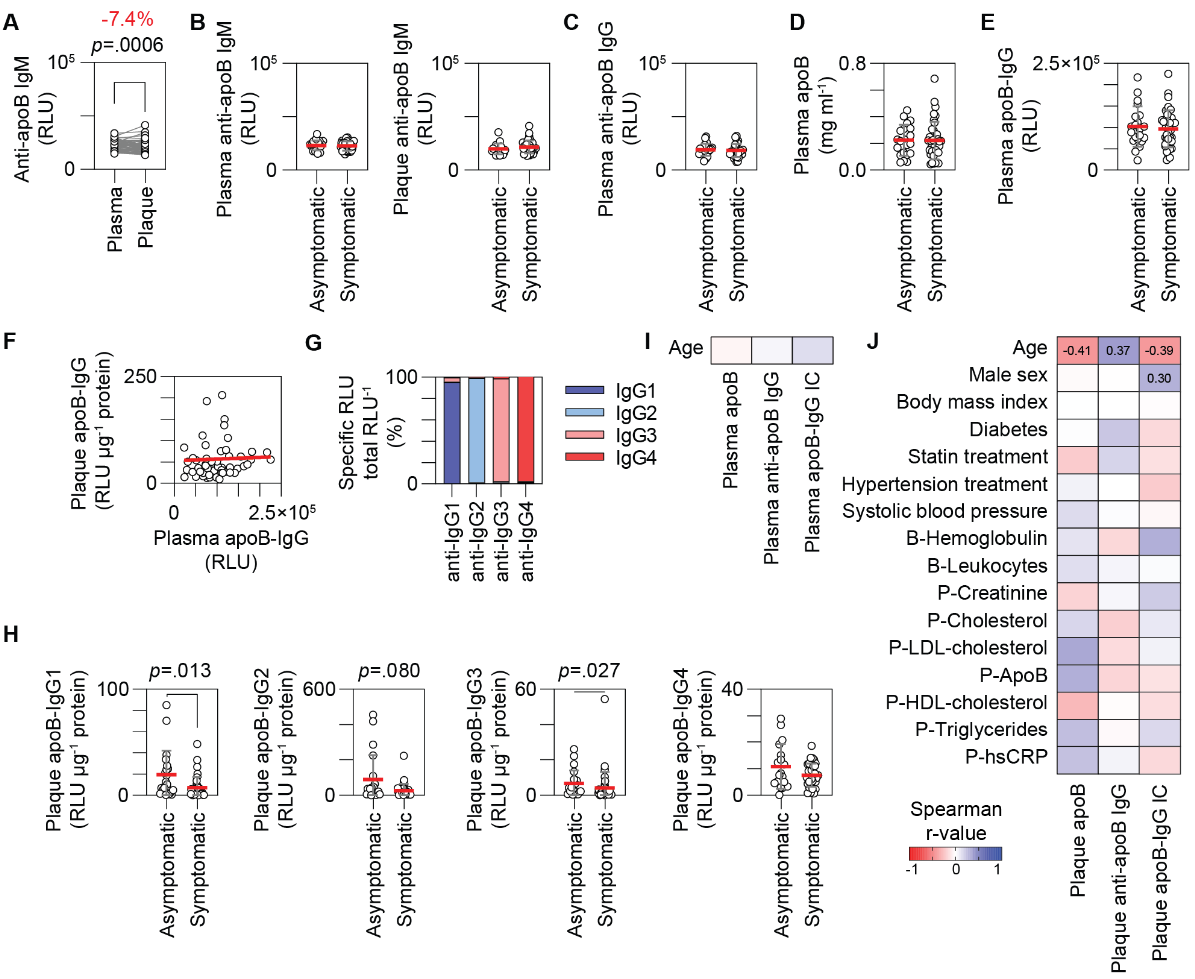


**Figure S5.** Associations between apoB-specific antibodies, clinical symptoms, and patient characteristics.
(**A**) Anti-apoB IgM levels in paired plasma and plaque samples normalized to 0.1 µg/ml total IgM (paired *t*-test).
(**B**) Anti-apoB IgM levels in plasma and plaques from asymptomatic versus symptomatic patients.
(**C**) Plasma anti-apoB IgG levels stratified by symptom status.
(**D**) Plasma apoB concentrations in asymptomatic versus symptomatic patients.
(**E**) Plasma levels of apoB-IgG immune complexes in asymptomatic and symptomatic patients.
(**F**) Relationship between apoB-IgG immune complexes in plasma and plaques (nonsignificant linear regression).
(**G**) Assay specificity for IgG isotypes: anti-IgG1 (95.1%), anti-IgG2 (97.8%), anti-IgG3 (96.5%), and anti-IgG4 (97.7%).
(**H**) Plaque levels of apoB-IgG1/IgG2/IgG3/IgG4 immune complexes in asymptomatic versus symptomatic patients, measured at 1:276 dilutions (IgG1-3) and 1:50 (IgG4) (Mann-Whitney test).
(**I**) Heatmap showing correlations between plasma apoB, anti-apoB IgG, and apoB-IgG immune complexes and patient age.
(**J**) Heatmap showing correlations between plaque apoB, anti-apoB IgG, and apoB-IgG immune complexes and clinical parameters; Spearman r-values displayed for significant associations ($p < .05$).
RLU, relative light units.

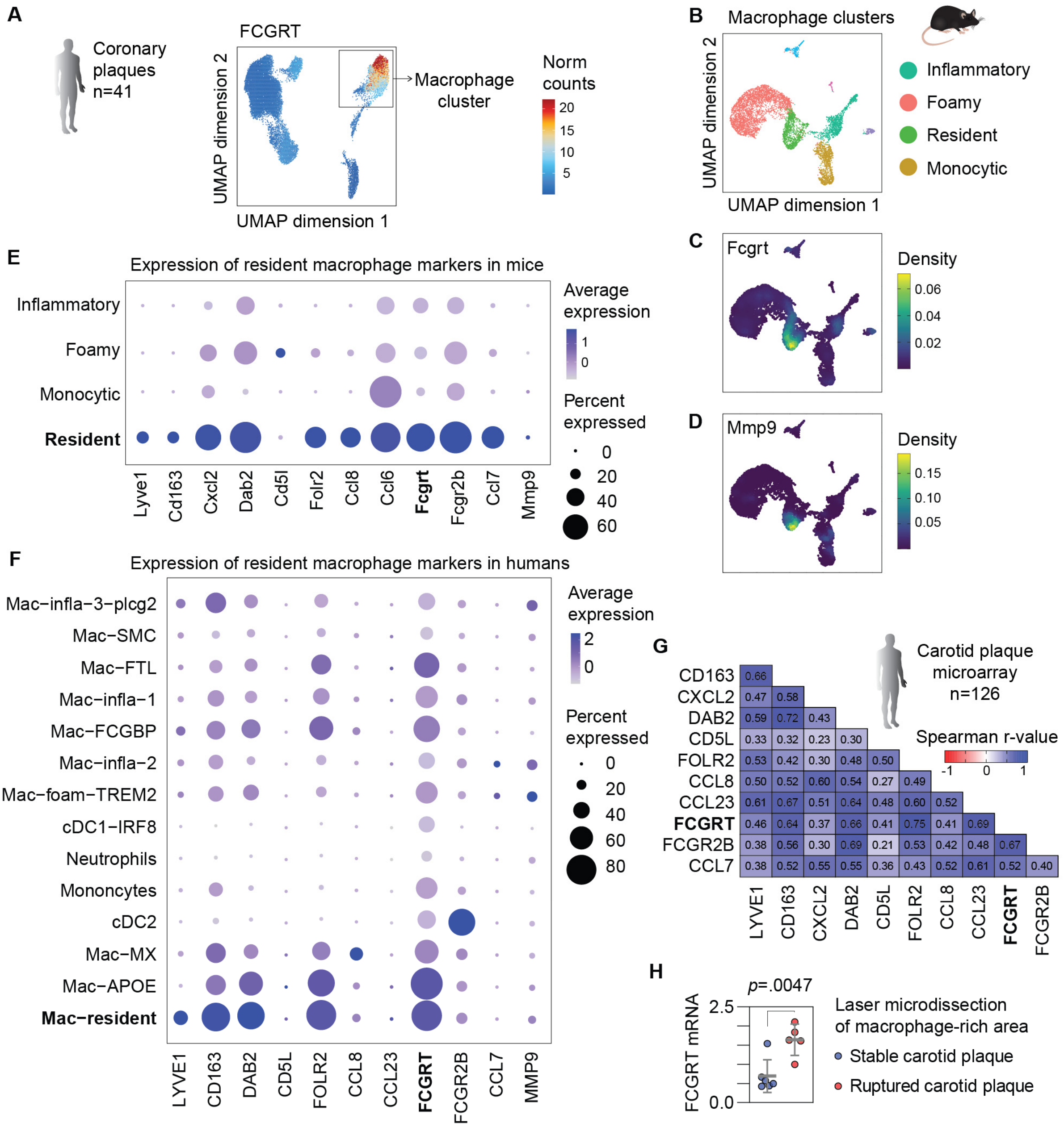


**Figure S6.** Expression of the neonatal Fc receptor in macrophage populations in atherosclerotic plaques.
(**A**) Single-cell assay for transposase-accessible chromatin sequencing dataset showing gene integration matrix for FCGRT in human coronary atherosclerotic plaque cells (Turner et al. *Nature Genetics* 2022); cluster annotations in Fig. S1F.
(**B**) Uniform manifold approximation and projection (UMAP) of macrophage clusters from mouse arterial tissue (GSE155513; Pan et al., *Circulation*, 2020). Arterial segments containing atherosclerotic lesions (ascending aorta, brachiocephalic artery, thoracic aorta) were isolated from *Ldlr*$^{-/-}$ mice on a Western-type diet, digested into single cells, and analyzed using 10x Chromium single-cell RNA-sequencing (14 samples, 30,251 cells). Macrophages were reclustered to identify four major populations.
(**C**) Feature plot of Fcgrt expression across macrophage clusters in the mouse dataset.
(**D**) Feature plot of Mmp9 expression across macrophage clusters in the mouse dataset.
(**E**) Dot plot showing resident-macrophage marker expression across macrophage clusters in the mouse dataset.
(**F**) Dot plot showing resident-macrophage marker expression across myeloid clusters in the integrated human single-cell RNA-sequencing dataset.

(**G**) Significant Spearman correlations among resident-macrophage markers in carotid plaque microarrays from 126 endarterectomy patients (GSE21545).
(**H**) FCGRT expression in laser-microdissected, macrophage-rich regions of ruptured versus stable human atherosclerotic plaques, based on microarray data with annotations derived from GPL570 (Lee et al., *Atherosclerosis*, 2012).

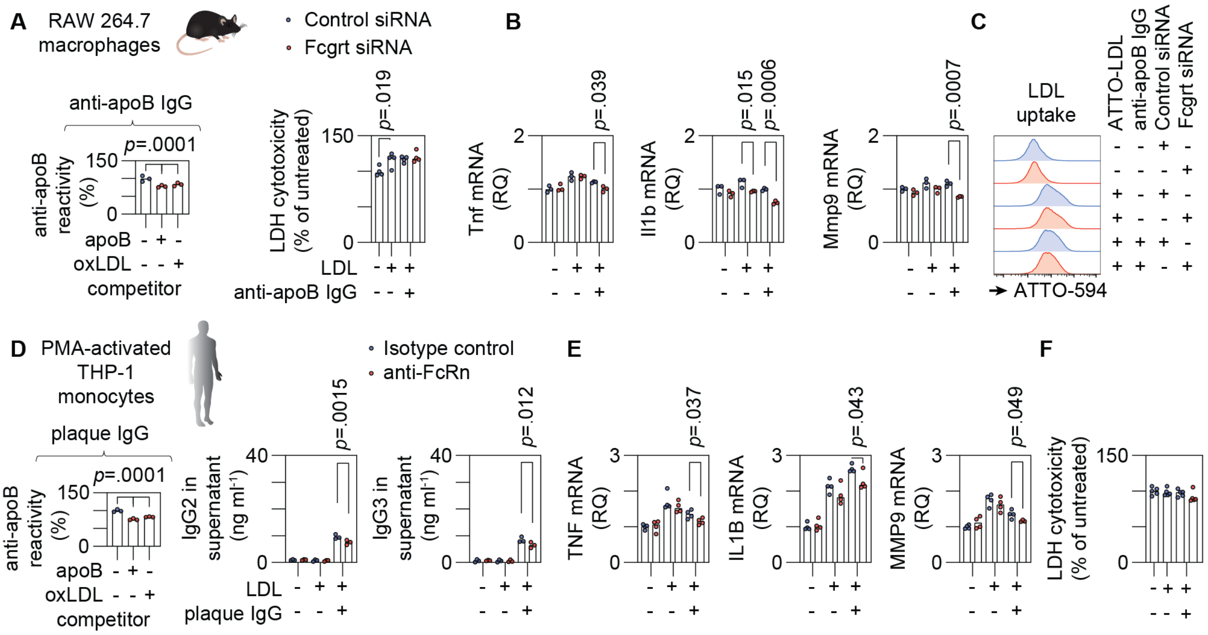


**Figure S7**. FcRn blockade suppresses IgG recycling and inflammatory activation in macrophages.
(**A**) Competitive ELISA confirming anti-apoB immunoreactivity in IgG isolated from mice receiving apoB-specific T cells (Gisterå et al., *Circulation*, 2018). Cross-reactivity with oxLDL was assessed using soluble competitors (10 µg/ml), reducing signal by -21.7% (apoB) and -16.4% (oxLDL). LDH cytotoxicity in RAW 264.7 macrophages (n = 4; one-way ANOVA with Holm-Šídák correction).
(**B**) mRNA expression changes in RAW 264.7 macrophages stimulated with immune complexes (n = 3; one-way ANOVA with Holm-Šídák correction).
(**C**) Representative histograms showing uptake of ATTO-594-labeled LDL by RAW 264.7 macrophages.
(**D**) Competitive ELISA demonstrating binding of plaque-derived IgG to apoB-coated plates. Soluble apoB or oxLDL (20 µg/ml) reduced binding by -24.5% (apoB) and -17.5% (oxLDL). IgG2 and IgG3 levels in supernatants of THP-1 macrophages stimulated with immune complexes (n = 3; one-way ANOVA with Holm-Šídák correction).
(**E**) mRNA expression in THP-1-derived macrophages stimulated with immune complexes (n = 4; unpaired *t*-test).
(**F**) LDH cytotoxicity in THP-1-derived macrophages (n = 5).
AU, arbitrary units; MFI, mean fluorescence index; RQ, relative quantification.

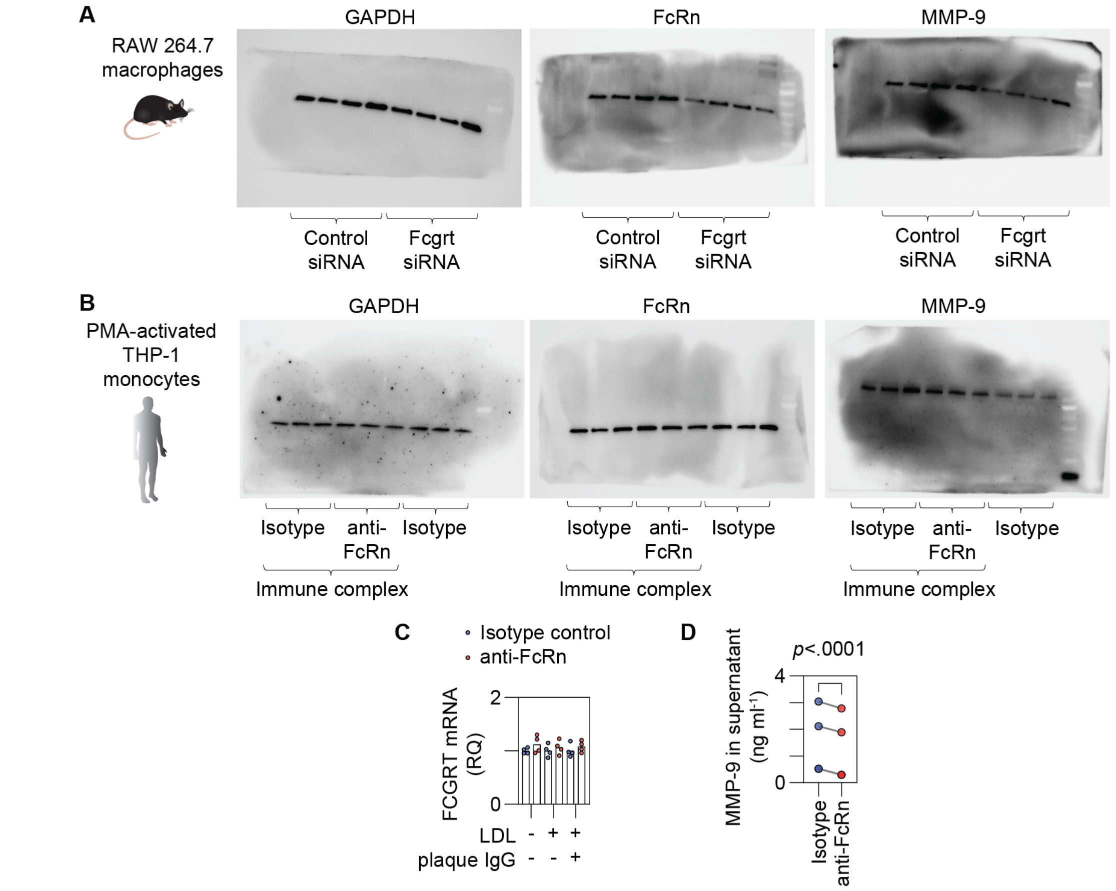


**Figure S8**. MMP-9 production in macrophages and human atherosclerotic plaques.
(**A**) Full, uncut immunoblots for GAPDH, FcRn, and MMP-9 in RAW 264.7 macrophages. Molecular weight markers: PageRuler Prestained Protein Ladder, 10-180 kDa (26616; ThermoFisher).
(**B**) Full, uncut immunoblots for GAPDH, FcRn, and MMP-9 in THP-1-derived macrophages using the same molecular weight ladder.
(**C**) FCGRT mRNA expression in THP-1-derived macrophages stimulated with immune complexes.
(**D**) Ex vivo stimulation of human carotid plaques for 24 hours, with MMP-9 quantified in culture supernatants (paired *t*-test; n = 4).

### Supplemental Tables

**Table S1.** Characteristics of patients with advanced carotid atherosclerosis included in the study.

| Characteristic | Immunostaining cohort<br>n=30 | Antibody extraction cohort<br>n=72 |
|---|---|---|
| **Age**<br>(years) | 71.0 ±7.9 | 69.6 ±9.1 |
| **Male sex**<br>(%) | 66.7% | 73.2% |
| **Symptoms within 6 months**<br>(%) | 89.7% | 64.8% |
| **Body weight**<br>(kg) | 75.7 ±14.1 | 76.9 ±12.9 |
| **Body Mass Index**<br>($kg/m^2$) | 25.8 ±4.3 | 26.2 ±3.8 |
| **Diabetes mellitus**<br>(%) | 13.3% | 19.7% |
| **Statin treatment**<br>(%) | 93.3% | 85.9% |
| **Hypertension treatment**<br>(%) | 60.0% | 85.7% |
| **Blood pressure**<br>(mmHg) | 142 ±22 / 77 ±10 | 153 ±18 / 80 ±8.9 |
| **Current smoker**<br>(%) | 26.9% | 16.9% |
| **Blood Hemoglobin**<br>(g/l) | 136 ±16 | 133 ±17 |
| **Blood Leukocytes**<br>($10^9$/l) | 7.4 ±1.9 | 7.4 ±2.0 |
| **Plasma Creatinine**<br>(µmol/l) | 75.4 ±12.5 | 91.2 ±21.4 |
| **Plasma Cholesterol**<br>(mmol/l) | 4.35 ±1.16 | 4.47 ±1.02 |
| **Plasma LDL-cholesterol**<br>(mmol/l) | 2.49 ±1.10 | 2.54 ±0.88 |
| **Plasma HDL-cholesterol**<br>(mmol/l) | 1.29 ±0.49 | 1.33 ±0.84 |
| **Plasma Triglycerides**<br>(mmol/l) | 1.20 ±0.58 | 1.59 ±1.09 |
| **Plasma hs-CRP**<br>(mg/l) | 1.10 [0.77-3.35] | 2.20 [0.93-4.35] |

hs-CRP=high-sensitivity C-reactive protein

**Table S2.** Antibodies used for immunohistochemistry, including clone, source, and working dilution.

| Target | Clone, RRID | Company, Catalog number | Species, conjugation | Concentration, Dilution |
|---|---|---|---|---|
| **Human IgG** | Polyclonal, AB_258513 | Sigma-Aldrich, B1140 | Goat, biotinylated | 3.3 mg/ml, 1:2,000 |
| **Human IgA** | 507, AB_109765 | Abcam, Ab124716 | Rabbit, unconjugated | 160 µg/ml, 1:400 |
| **Human IgM** | Polyclonal, AB_258514 | Sigma-Aldrich, B1265 | Goat, biotinylated | 2.5 mg/ml, 1:2,000 |
| **Human IgE** | Polyclonal, AB_374853 | GeneTex, GTX73360 | Rabbit, unconjugated | 5.9 mg/ml, 1:200 |
| **Isotype control** | Polyclonal, AB_3675839 | Abcam, Ab37376 | Goat, biotinylated | 0.5 mg/ml, 1:2,000 |
| **Goat IgG** | Polyclonal, AB_2336123 | Vector Labs, BA9500 | Horse, biotinylated | 1.5 mg/ml, 1:200 |
| **Mouse IgG** | Polyclonal, AB_2313581 | Vector Labs, BA2000 | Horse, biotinylated | 1.5 mg/ml, 1:200 |
| **Rabbit IgG** | Polyclonal, AB_2313606 | Vector Labs, BA1000 | Goat, biotinylated | 1.5 mg/ml, 1:200 |
| **Human CD79A** | JCB117, AB_1158199 | Sigma-Aldrich, 179M-94 | Mouse, unconjugated | 51 µg/ml, 1:300 |
| **Human CD3ε** | PS1, AB_3552706 | BioCare Medical, PM110AA | Mouse, unconjugated | 3.2 µg/ml, prediluted |
| **Isotype control** | G155-178, AB_394869 | BD Biosciences, 553455 | Mouse, biotinylated | 0.5 mg/ml, 1:200 |
| **Malondialdehyde** | LR04 IgM, N/A | Binder Lab, Vienna N/A | Mouse, unconjugated | 2.5 mg/ml, 1:2,500 |
| **NLRP3** | Cryo-2, AB_2490202 | AdipoGen, AG-20B-0014 | Mouse, unconjugated | 1.0 mg/ml, 1:200 |
| **Mouse IgM** | Polyclonal, AB_2336183 | Vector Labs, BA2020 | Goat, biotinylated | 0.5 mg/ml, 1:200 |
| **Human CD68** | PG-M1, AB_2074844 | Dako, M0876 | Mouse, unconjugated | 40 µg/ml, 1:200 |
| **Human α-actin** | 1A4, AB_2223500 | Dako, M0851 | Mouse, unconjugated | 71 µg/ml, 1:400 |
| **Human FcRn** | CL3638, AB_2637274 | Thermo Fisher, MA5-24659 | Mouse, unconjugated | 1 mg/ml, 1:400 |
| **CD163** | Polyclonal K-18, AB_2291274 | Santa Cruz, sc-18796 | Goat, unconjugated | 0.2 mg/ml, 1:300 |
| **Mouse IgG** | Polyclonal, AB_2336412 | Vector Labs, DI-2594 | Horse, Dylight 594 | 1.5 mg/ml, 1:200 |
| **Goat IgG** | Polyclonal, AB_2336400 | Vector Labs, DI-3088 | Horse, Dylight 488 | 1.5 mg/ml, 1:200 |

RRID=Research Resource Identifiers

**Table S3.** Antibodies used for immunoassays, including capture, detection, and isotype-specific reagents.

| Target | Clone, RRID | Company, Catalog number | Species, conjugation | Concentration, Dilution |
|---|---|---|---|---|
| **Human IgG** | MT145, AB_10697677 | Mabtech, 3850-1-250 | Mouse, unconjugated | 0.5 mg/ml, 1:250 |
| **Human IgA** | MT57, AB_10736405 | Mabtech, 3860-3-250 | Rat, unconjugated | 0.5 mg/ml, 1:250 |
| **Human IgM** | G20-127, AB_396115 | BD Biosciences, 555780 | Mouse, unconjugated | 0.5 mg/ml, 1:250 |
| **Human IgE** | 107, AB_907287 | Mabtech, 3810-3-250 | Mouse, unconjugated | 1 mg/ml, 1:500 |
| **Human IgG** | MT78, AB_10697678 | Mabtech, 3850-9A | Mouse, alkaline phosphatase | 0.64 mg/ml 1:1,000 |
| **Human IgM** | Polyclonal, AB_258474 | Sigma-Aldrich, A9794 | Goat, alkaline phosphatase | 6.9 mg/ml, 1:2,000 |
| **Human IgG1** | Polyclonal, AB_3675840 | Southern Biotech, 9050-04 | Mouse, alkaline phosphatase | 0.4 mg/ml, 1:2,000 |
| **Human IgG2** | HP6014, AB_2796645 | Southern Biotech, 9080-04 | Mouse, alkaline phosphatase | 1.7 mg/ml, 1:2,000 |
| **Human IgG3** | HP6050, AB_2687998 | Southern Biotech, 9210-04 | Mouse, alkaline phosphatase | 0.86 mg/ml, 1:2,000 |
| **Human IgG4** | HP6023, AB_2796683 | Southern Biotech, 9190-04 | Mouse, alkaline phosphatase | 1.85 mg/ml, 1:2,000 |
| **Human IgG1 isotype control** | Myeloma serum, AB_2794079 | Southern Biotech, 0151K-01 | Human, unconjugated | 0.5 mg/ml |
| **Human IgG2 isotype control** | Myeloma serum, AB_2715459 | BioXCell, BE0301 | Human, unconjugated | 6.63 mg/ml |
| **Human IgG3 isotype control** | Myeloma serum, AB_2794088 | Southern Biotech, 0153L-01 | Human, unconjugated | 0.5 mg/ml |
| **Human IgG4 isotype control** | Monoclonal, AB_3739870 | MedChemExpress, HY-P99003 | Human, unconjugated | 5.31 mg/ml |
| **Human FcRn** | Rozanolixizumab, AB_3695464 | MedChemExpress, HY-P9979 | Human, unconjugated | 5.34 mg/ml |
| **Human apoB** | Polyclonal, AB_10816186 | Biosynth, 20-AG40S | Goat, unconjugated | 4.78 mg/ml, 1:500 |
| **Mouse IgG** | Polyclonal, AB_2794303 | Southern Biotech, 1031-01 | Goat, unconjugated | 1.0 mg/ml, 1:500 |
| **Mouse IgG** | Polyclonal, AB_2794293 | Southern Biotech, 1030-04 | Goat, alkaline phosphatase | 1.0 mg/ml, 1:5,000 |
| **FcRn** | Polyclonal, AB_2812353 | Thermo Fisher, PA5-97738 | Rabbit, unconjugated | 1.0 mg/ml 1:1,000 |
| **Human MMP-9** | Polyclonal, AB_2880837 | Proteintech, 27306-1-AP | Rabbit, unconjugated | 350 µg/ml 1:1,000 |

| | | | | |
|---|---|---|---|---|
| **Mouse MMP-9** | 5A1,<br>AB_3086555 | Proteintech,<br>82854-1-RR | Rabbit,<br>unconjugated | 1.0 mg/ml<br>1:2,000 |
| **GAPDH** | EPR16891,<br>AB_2630358 | Abcam,<br>ab181602 | Rabbit,<br>unconjugated | 1.1 mg/ml<br>1:2,000 |
| **Mouse IgG** | Polyclonal,<br>AB_2636929 | Dako,<br>P0260 | Rabbit, horseradish<br>peroxidase | 0.25 mg/ml<br>1:2,000 |
| **Rabbit IgG** | Polyclonal,<br>AB_2617138 | Dako,<br>P0448 | Goat, horseradish<br>peroxidase | 0.25 mg/ml<br>1:2,000 |

RRID=Research Resource Identifiers

**Table S4.** Assay-on-demand probes and siRNA reagents used for real-time PCR and gene-silencing experiments.

| Target | Reference number |
|---|---|
| Hprt | Mm01545399_m1 |
| Fcgrt | Mm01205451_m1 |
| Il1b | Mm00434228_m1 |
| Mmp9 | Mm00442991_m1 |
| Mouse Fcgrt siRNA | 4390771 |
| Silencer Select Negative Control No. 1 siRNA | 4390843 |
| PPIA | Hs99999904_m1 |
| FCGRT | Hs00175415_m1 |
| IL1B | Hs01555410_m1 |
| MMP9 | Hs00234579_m1 |
| TNF | Hs00174128_m1 |
